# TomoSwin3D: a Swin3D Transformer for the Identification and Classification of Macromolecules in 3D Cryo-ET Tomograms

**DOI:** 10.64898/2026.04.17.719219

**Authors:** Ashwin Dhakal, Rajan Gyawali, Jianlin Cheng

**Affiliations:** Department of Electrical Engineering and Computer Science, University of Missouri, Columbia, MO 65211, USA; NextGen Precision Health, University of Missouri, Columbia, MO 65211, USA

## Abstract

Cryo-electron tomography (cryo-ET) enables in situ three-dimensional visualization of many protein complexes and other macromolecular assemblies such as ribosomes in cells, yet automated macromolecule particle identification in 3D cryo-ET tomograms remains a major bottleneck due to dose-limited low signal-to-noise ratios, missing-wedge artifacts, and densely crowded cellular backgrounds. We present TomoSwin3D, an end-to-end three-dimensional (3D) macromolecule particle identification and classification pipeline centered on a Swin Transformer-based U-Net that performs particle identification and classification and outputs particle centroid coordinates. TomoSwin3D leverages a multi-channel input representation that augments raw tomogram densities with complementary 3D feature maps capturing edge strength (Sobel gradients), local contrast enhancement (morphological top-hat), and multiscale blob responses (Difference-of-Gaussians), improving detectability of small and low-contrast targets. To better preserve particle geometry and avoid hand-crafted shape assumptions, it adopts occupancy-preserving supervision that directly uses available 3D instance masks rather than heuristic Gaussian/spherical labels and applies scalable patch-wise inference followed by lightweight post-processing (connected-component analysis, size filtering, centroid extraction) for robust centroid coordinate extraction. Across diverse simulated and experimental cryo-ET tomogram benchmarks including SHREC 2021 and 2020 test datasets, EMPIAR dataset, and Cryo-ET data portal dataset, TomoSwin3D achieves strong and consistent performance in detecting proteins and other particles, outperforming existing methods, with a pronounced advantage in picking hard, small protein particles. These results establish TomoSwin3D as a scalable and accurate solution for high-throughput cryo-ET macromolecule particle picking and downstream subtomogram averaging.

## 1. Introduction

### 1.1 The significance of cryo-electron tomography (cryo-ET)

Cryo-ET has emerged as a powerful imaging technology capable of producing 3D visualizations of macromolecular complexes such as protein assemblies in their native cellular environment. Unlike traditional structural biology methods that require protein purification, cryo-ET preserves samples in a vitrified state, allowing researchers to explore the molecular details of the cell at sub-nanometer to near-atomic resolutions ^1^. This capability is the foundation of visual proteomics, a field aimed at mapping the spatial distribution, interactions, and conformational states of the entire cellular proteome to create a comprehensive “atlas” of the cell.

Despite its potential, individual tomograms are fundamentally limited by physical constraints. To minimize radiation damage to sensitive biological samples, a limited electron dose is applied, resulting in an extremely low signal-to-noise ratio (SNR), typically below 0.1 ^2^. Furthermore, because the sample cannot be tilted through a full 360-degree range, the resulting data suffers from a “missing wedge” artifact in Fourier space, which leads to anisotropic resolution and the delocalization of structural densities along the Z-axis. To overcome these limitations, researchers utilize subtomogram averaging (STA) ^3^, a process that identifies, aligns, and integrates thousands of noisy copies of a specific macromolecule to improve the signal and resolve high-resolution structural details ^4^.

### 1.2 The “particle identification and classification” bottleneck

The first and most critical step in the STA pipeline for proteome analysis is protein particle identification (also called particle picking), which involves localizing the 3D coordinates and orientations of target proteins within crowded cellular volumes. This task has become the primary computational and logistical bottleneck in structural biology ^5 6^. Manual picking by experts is subjective, prone to bias, and prohibitively labor-intensive, often requiring lengthy time to annotate a single tomogram. The challenge is exacerbated by the densely crowded environment of the cell, where molecular weight heterogeneity and structural similarity make it difficult to distinguish specific targets from background noise, organelles, and membranes^2^. This difficulty is particularly acute for small macromolecules (<200 kDa), which offer low contrast and are easily mistaken for imaging artifacts.

### 1.3 Evolution and limitations of existing particle picking methods

The field of protein and other macromolecule particle picking in cryo-ET has evolved from manually intensive geometric methods toward automated deep learning (DL) architectures that prioritize speed and annotation efficiency. Conventional template-based methods ^7–10^ have historically served as the standard, primarily utilizing template matching (TM), which employs 3D cross-correlation to scan tomograms using reference models from databases like the Protein Data Bank (PDB) ^11^. While TM provides essential orientation information for STA, it is computationally expensive, taking hours per tomogram, and is highly susceptible to reference bias and false positives in low SNR cellular environments. To address the need for reference-free detection, Difference of Gaussian (DoG) filtering was introduced to identify structural edges, though it struggles with the high noise levels typical of in situ data. Interactive geometric tools such as Dynamo ^12^, Blik ^13^, and MPicker ^14^ further streamlined picking by allowing users to model filaments or flatten 3D membranes into 2D maps, yet these remain labor-intensive and require significant human intervention.

The introduction of supervised deep learning marked a paradigm shift, as models began learning complex structural features directly from noisy data. Early DL frameworks, such as DeepFinder ^15^ and EMAN2 ^16^, utilized 3D U-Net architectures to perform voxel-wise semantic segmentation, classifying cellular content into discrete classes like “ribosome” or “membrane”. While these methods significantly outperform TM in accuracy, their primary limitation is the annotation burden, often requiring thousands of manual voxel-level labels for training. To improve inference speed, object detection frameworks like VP-Detector ^17^ and PickYOLO ^18^ were developed; these treat picking as a coordinate prediction task, bypassing expensive post-segmentation clustering. Advancements in this era also produced specialized architectures, including 3D-UCaps ^19^ for rotation invariance, FSPicker ^20^ for dual-stream multi-scale attention, and TomoCPT ^21^, which utilizes transformer-based SwinUNETR ^22^ modules to predict particle centroids using Gaussian labels rather than binary masks.

The current frontier of cryo-ET research focuses on generalization and annotation efficiency to enable high-throughput structural discovery. Recent “universal” pickers like TomoTwin ^23^ utilize deep metric learning to map subvolumes into a high-dimensional embedding space, allowing the identification of novel proteins without retraining. Similarly, ProPicker ^24^ introduces a promptable segmentation architecture that selectively detects target proteins based on a single input prompt, while CryoSAM ^25^ leverages general-domain foundation models to segment particles slice-by-slice. To further reduce human labor, researchers have explored weakly supervised and positive-unlabeled (PU) learning; for instance, TomoPicker ^26^ can train effectively on a minuscule fraction of a dataset, and RobPicker ^27^ employs a meta-learning framework with label-correction and reweighting networks to handle noisy or imbalanced training data. Fully unsupervised methods, such as PickET ^28^, represent a different approach, using Gabor features and de novo clustering to segment cellular content without any structural templates or labeled training data.

### 1.4 The proposed methodology

Despite rapid progress in cryo-ET protein and other particle picking, existing approaches still struggle in the setting that matters most for high-throughput in situ structural biology: small, low-contrast particles embedded in crowded cellular background, under dose-limited low SNR and missing-wedge–induced distortions. Template matching is expensive and prone to false positives in cellular environments, while supervised deep learning methods often require dense voxel-level labels or large, carefully curated training sets. In practice, available annotations are frequently sparse, inconsistent, and noisy, and performance can drop when models are applied across tomograms with different acquisition conditions or sample types. These limitations make particle picking a persistent bottleneck for STA, especially when scaling to thousands to millions of particles where both precision (to avoid downstream contamination) and recall (to maximize usable particles) are critical.

To address this gap, we propose a deep learning–based 3D particle picker designed for robust coordinate prediction in cryo-ET tomograms, with a focus on improving detection of small and difficult targets while remaining practical for large-scale datasets. Our approach uses a 3D Swin Transformer ^29^–based U-Net ^30^ architecture (Swin3D U-Net) that combines hierarchical, windowed self-attention with a multi-resolution encoder–decoder design. The model operates on multi-channel tomogram representations, enabling it to integrate complementary cues from the raw tomogram reconstruction and derived feature channels, and to better separate true particle signal from structured cellular background. The network performs multiclass particle identification that is converted into particle coordinates through lightweight post-processing, enabling efficient inference on large tomogram volumes.

In summary, TomoSwin3D is designed to (i) increase picking accuracy for small/low-contrast particles, (ii) remain robust under low SNR and missing-wedge artifacts with high F1 score, and (iii) support scalable inference suitable for high-throughput STA pipelines. We demonstrate that this architecture yields improved detection performance compared to existing methods as detailed in the results section, enabling higher-quality particle sets for downstream subtomogram averaging and analysis.

## 2. Results

### 2.1 Problem definition and workflow

TomoSwin3D is an end-to-end 3D particle-picking framework that takes as input a raw cryo-electron tomogram and outputs coordinate files of macromolecules and the annotated 3D volume (**Figure 1**). Briefly, each tomogram (**Figure 1A**) is first standardized by resampling to a common voxel size and applying percentile-based intensity normalization with clipping to map densities into a consistent ([0,1]) range (**Figure 1B**). The normalized tomograms are then split into 3D sub-volumes (**Figure 1C**) using a uniform context-preserving window (64×64×64) to enable training and inference on large tomograms. Input subvolumes (grids) are augmented into four-channel tensors by stacking normalized sub-tomogram with 3D feature maps: multi-scale difference-of-Gaussians (DoG) for blob enhancement, Sobel gradient magnitude for edge detection, and multi-scale top-hat transforms for local contrast improvement (**Figure 1D**).

**Figure 1:**
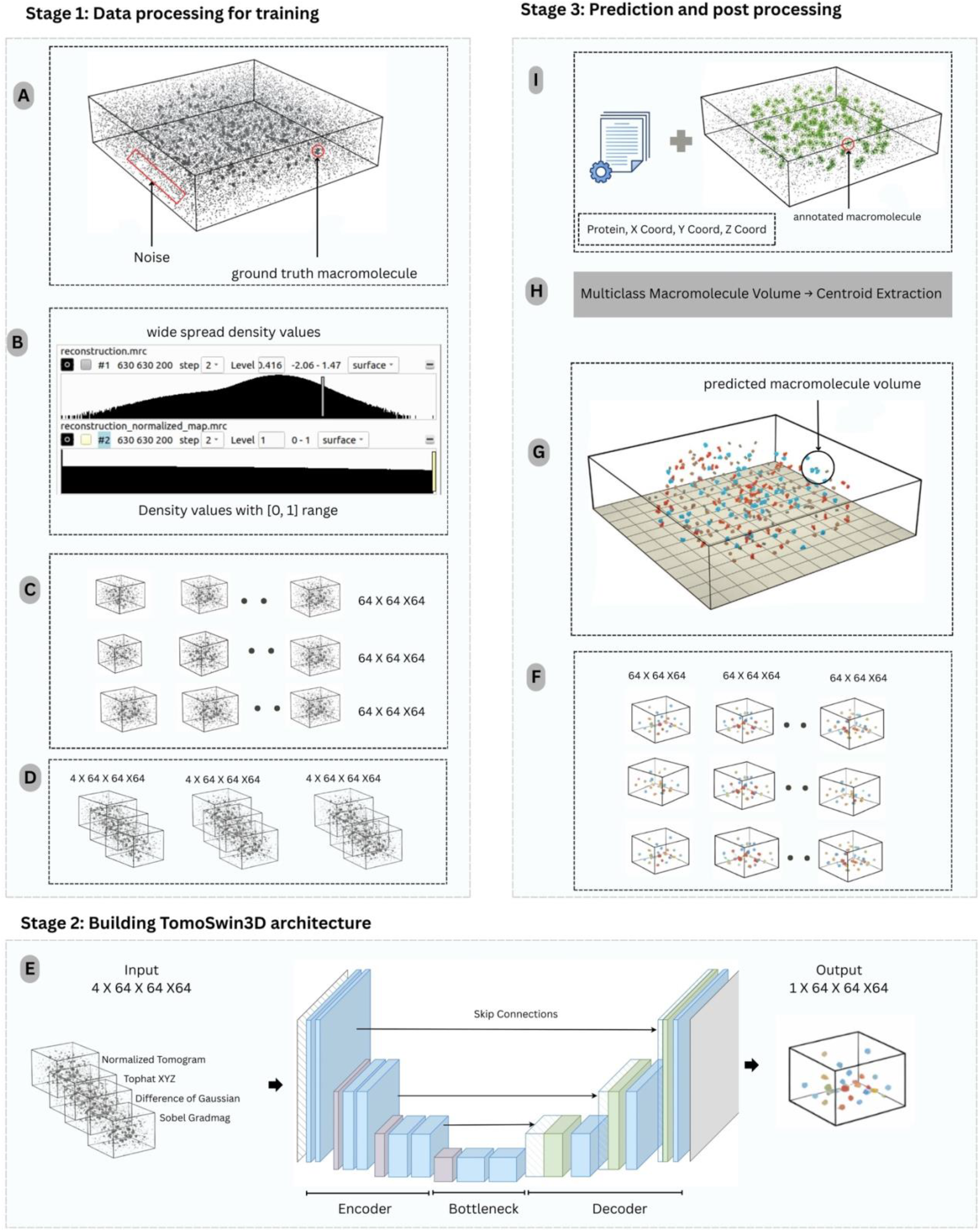
Overall workflow of TomoSwin3D. **(A)** Raw input tomogram containing target macromolecules in strong background noise. **(B)** Intensity normalization: original voxel-density distribution (top) and normalized values scaled to ([0,1]) (bottom). **(C)** The normalized tomogram partitioned into sub-volumes (grids) of size (64×64×64). **(D)** A multi-channel input tensor constructed by stacking the normalized sub-tomogram with feature-enhancement maps. **(E)** representative TomoSwin3D architecture (3D U-Net with attention-based encoder–decoder) takes the 4-channel input and produces a voxel-wise prediction volume. **(F)** Generated (64×64×64) output grids. **(G)** Grid-level predictions stitched to reconstruct a single multiclass macromolecule volume matching the original tomogram dimensions. **(H)** Connected-component analysis and centroid extraction applied to each predicted macromolecule region to obtain particle centers. **(I)** Final output consisting of macromolecule centroid coordinates (x,y,z) and the corresponding 3D annotated volume for downstream analysis and visualization.

TomoSwin3D employs a 3D Swin Transformer integrated into a U-Net-style encoder–decoder. The encoder progressively downsamples the input through learnable patch merging while applying window-based self-attention with shifted windows to capture both local and cross-window context efficiently in 3D; the decoder mirrors this process with learnable upsampling and skip connections to recover fine spatial detail, followed by a final prediction head that outputs per-voxel class logits for binary or multiclass segmentation (**Figure 1E**).

For inference, test tomograms are processed patch-wise by partitioning each input tomogram into fixed-size subvolumes (grids). The network outputs voxel-wise multiclass probability maps for every grid, which are then thresholded to suppress low-confidence labels (**Figure 1F**). Grid-level predictions are subsequently stitched into a tomogram-sized volume which outputs the same size as the input tomogram (**Figure 1G**). Post-processing converts the dense semantic segmentation into sparse class-aware particle coordinates. Specifically, for each predicted class, a 3D connected-component analysis is performed using 18-connectivity to identify discrete particle blobs. Components smaller than a minimum size threshold are removed to reduce spurious detections arising from noise and reconstruction artifacts. For each retained component, a particle coordinate is computed as the centroid (center-of-mass) of the component voxels (**Figure 1H & Figure 1I**). Predicted and ground truth coordinates are matched using one-to-one Hungarian assignment under a distance cutoff, and detection performance is quantified using precision, recall, and F1 score, and other metrics as reported in the Results Section.

### 2.2 Performance of TomoSwin3D on SHREC 2021 synthetic test data

On the standard test data of 2021 3D Shape Retrieval Challenge (SHREC) competition ^31^, TomoSwin3D achieves the highest overall accuracy among all compared methods, with a mean F1-score of 0.861 across 13 macromolecule targets, belonging to 12 protein classes and a reference fiducial (see particle shape, size, and weights visualized in **Figure 2A**).

**Figure 2:**
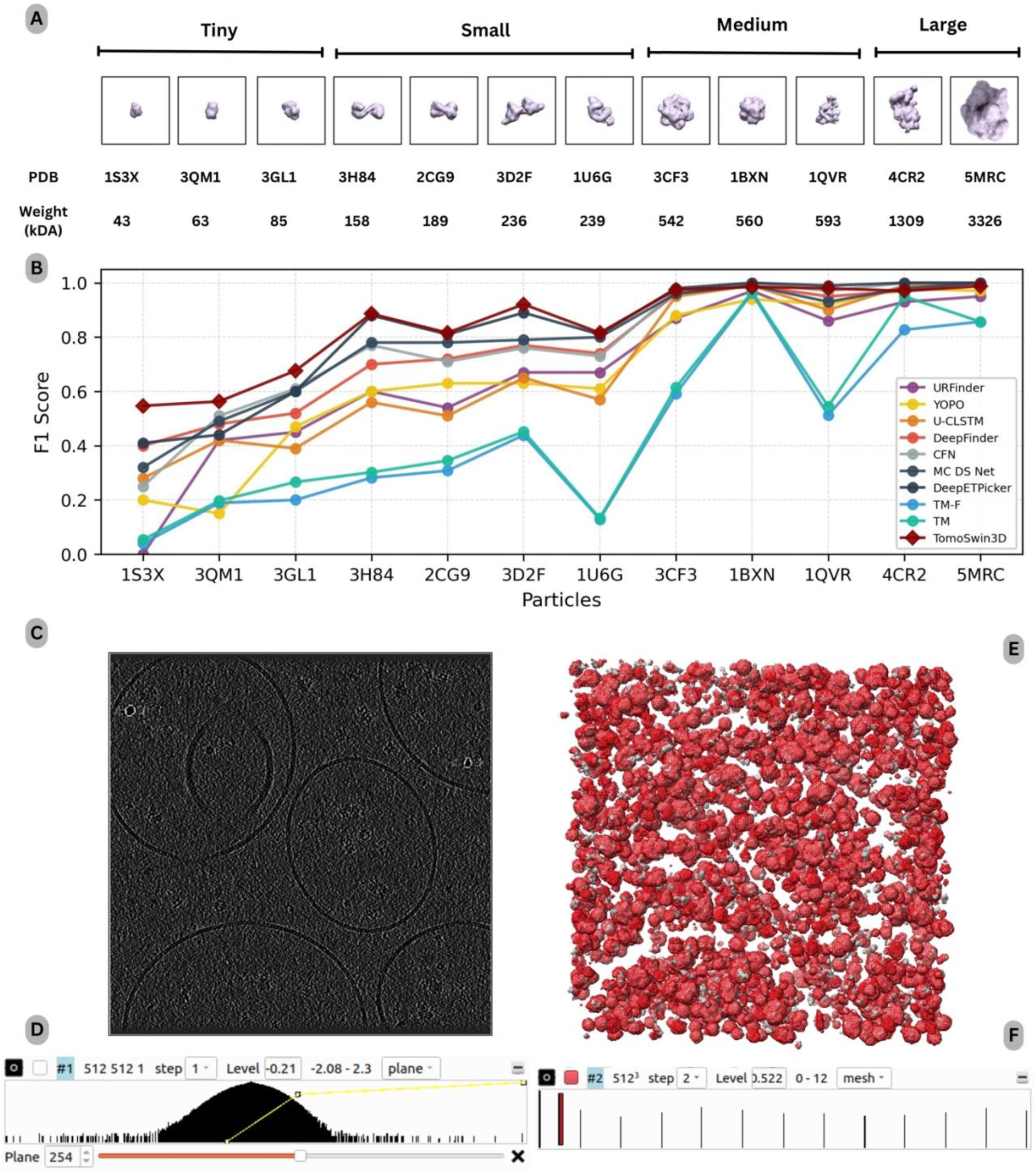
Performance analysis on the SHREC 2021 (synthetic) dataset. **(A)** Overview of the SHREC 2021 test proteins grouped by size/weight (kDa) and corresponding PDB identifiers. **(B)** Per-class F1-score comparison across competing methods for multiclass particle detection. **(C)** Representative input tomogram slice at (z=254), illustrating the challenging low-SNR background and crowded scene. **(D)** Voxel-intensity histogram of the raw tomogram, highlighting the broad dynamic range prior to normalization. **(E)** 3D qualitative comparison between ground-truth macromolecules (grey) and TomoSwin3D predictions (red mesh) in the reconstructed volume. **(F)** Volume-viewer rendering (UCSF Chimera) showing the spatial distribution of predicted multiclass macromolecules across the tomogram.

As shown in **Table 1**, there is an absolute improvement of +0.031 F1-score over the state-of-the-art deep-learning method (DeepETPicker ^32^, 0.830) and a much larger margin over other learning-based methods such as MC DS Net (0.800), CFNPicker ^33^ (0.790), and DeepFinder ^15^ (0.780). In contrast, the F1-score of classical template matching baselines is substantially lower (TM: 0.51; TM-F: 0.48), highlighting the difficulty of the SHREC 2021 cellular tomograms for correlation-based scanning and the benefit of learned, data-adaptive feature representations.

**Table 1:** Multiclass particle-detection performance comparison of different methods on the SHREC 2021 (synthetic) test set.

| Method/Proteins | 1S3X | 3QM1 | 3GL1 | 3H84 | 2CG9 | 3D2F | 1U6G | 3CF3 | 1BXN | 1QVR | 4CR2 | 5MRC | Fiducial | Mean F1 |
| --- | --- | --- | --- | --- | --- | --- | --- | --- | --- | --- | --- | --- | --- | --- |
| **URFinder** | 0 | 0.42 | 0.45 | 0.6 | 0.54 | 0.67 | 0.67 | 0.87 | 0.97 | 0.86 | 0.93 | 0.95 | 0.43 | 0.64 |
| **YOPO** | 0.2 | 0.15 | 0.47 | 0.6 | 0.63 | 0.63 | 0.61 | 0.88 | 0.94 | 0.92 | 0.98 | 0.97 | 0.95 | 0.69 |
| **U-CLSTM** | 0.28 | 0.42 | 0.39 | 0.56 | 0.51 | 0.65 | 0.57 | 0.95 | 0.99 | 0.9 | 0.99 | **1** | **1** | 0.71 |
| **DeepFinder** | 0.4 | 0.48 | 0.52 | 0.7 | 0.72 | 0.77 | 0.74 | 0.96 | 0.99 | 0.95 | 0.97 | **1** | **1** | 0.78 |
| **CFNPicker** | 0.25 | 0.51 | 0.61 | 0.77 | 0.71 | 0.76 | 0.73 | 0.97 | **1** | 0.97 | **1** | **1** | **1** | 0.79 |
| **MC DS Net** | 0.32 | 0.49 | 0.6 | 0.78 | 0.78 | 0.79 | 0.8 | 0.96 | 0.99 | 0.93 | 0.98 | **1** | **1** | 0.8 |
| **DeepETPicker** | 0.41 | 0.44 | 0.6 | 0.88 | 0.81 | 0.89 | 0.81 | **0.98** | **1** | 0.99 | **1** | **1** | **1** | 0.83 |
| **TM-F** | 0.04 | 0.189 | 0.2 | 0.282 | 0.308 | 0.439 | 0.129 | 0.592 | 0.962 | 0.513 | 0.827 | 0.857 | 0.9 | 0.48 |
| **TM** | 0.054 | 0.197 | 0.266 | 0.302 | 0.345 | 0.452 | 0.133 | 0.615 | 0.966 | 0.545 | 0.95 | 0.857 | 0.9 | 0.51 |
| **TomoSwin3D (ours)** | **0.547** | **0.563** | **0.676** | **0.886** | **0.816** | **0.921** | **0.816** | 0.975 | 0.985 | 0.98 | 0.97 | 0.988 | **1** | **0.861** |

A key observation from the class-wise results is that TomoSwin3D’s advantage is concentrated on the most challenging small-particle classes (e.g., 1S3X, 3QM1, 3GL1) where competing methods show large performance drops, as shown in **Figure 2B**. Specifically, TomoSwin3D attains the best F1-scores on 7 of the 12 protein targets: 1S3X (0.547), 3QM1 (0.563), 3GL1 (0.676), 3H84 (0.886), 2CG9 (0.816), 3D2F (0.921), and 1U6G (0.816), and ties for best on fiducials (1.000). Detailed metrics are provided in **Supplementary Table S1**. These gains are particularly pronounced for classes that appear to be systematically difficult for other approaches (e.g., 1S3X and 3QM1, where most baselines remain below ∼0.5), suggesting improved robustness to low contrast, and ambiguity in local context. For example, on 1S3X, TomoSwin3D improves from the next-best F1 of 0.41 (DeepETPicker) to 0.547 (absolute +0.137), indicating a substantial reduction in combined false positives and false negatives for this hard class (test tomogram and its raw density values shown in **Figure 2C&D** and output predictions and multiclass values shown in **Figure 2E&F)**.

For several larger or more distinctive targets: such as 1BXN, 4CR2, and 5MRC (details of macromolecules included in **Supplementary Table S2, S3 & S4)**, multiple deep learning methods in TomoSwin3D approach the metric ceiling (often F1 ≈ 1.0). In these cases, TomoSwin3D remains highly competitive (0.985 - 0.988), but does not always exceed the best reported scores, which is expected when performance is already near saturation and residual differences may slightly fluctuate with borderline detections or annotation tolerance effects. Importantly, despite these near-ceiling patterns, TomoSwin3D maintains strong consistency across all targets, with F1 ≥ 0.816 on 10 of 13 targets, and no catastrophic failure cases (in contrast to the template-matching baselines, which exhibit very low F1 for multiple protein classes).

**Table 2** summarizes binary detection performance on the SHREC 2021 test data, where all protein particles are treated as a single foreground class. Under this setting, TomoSwin3D attains the best overall performance, achieving an F1-score of 0.947 with precision = 0.990 and recall = 0.907. This result improves upon the strongest competing deep-learning baseline, DeepETPicker (F1 = 0.939; precision = 0.958; recall = 0.921), and substantially exceeds earlier CNN/RNN-based approaches (e.g., DeepFinder: 0.868, MC DS Net: 0.850, U-CLSTM: 0.827, CFNPicker: 0.818). Template matching remains markedly inferior (TM-F: 0.576; TM: 0.372), consistent with its known sensitivity to low SNR and structural heterogeneity in crowded cellular tomograms.

**Table 2:** Binary particle detection performance comparison of existing methods on the SHREC 2021 (synthetic) test set (RR: reported results i.e., number of particles detected by each method; TP: true positives, representing correctly identified unique particles; FP: false positives, indicating detections of particles that do not exist; FN: false negatives, referring to actual particles that were missed; MH: multiple hits, where a single true particle is detected more than once; AD: mean Euclidean distance (in voxels) between predicted and actual particle centers; Recall: ratio of correctly identified unique particles to the total number of true particles in the test tomogram; Precision: ratio of correctly identified unique particles to the total reported results (RR); F1 Score: harmonic mean of precision and recall).

| Method | RR | TP | FP | FN | MH | AD | Recall ↑ | Precision ↑ | F1 Score ↑ |
| --- | --- | --- | --- | --- | --- | --- | --- | --- | --- |
| **URFinder** | 1969 | 1298 | 377 | 267 | 149 | 1.84 | 0.826 | 0.659 | 0.733 |
| **YOPO** | 1627 | 1224 | 232 | 341 | 14 | 1.66 | 0.72 | 0.752 | 0.765 |
| **CFNPicker** | 1765 | 1364 | 239 | 201 | 20 | 1.52 | 0.868 | 0.773 | 0.818 |
| **U-CLSTM** | 1460 | 1253 | 49 | 312 | 44 | 2.13 | 0.798 | 0.858 | 0.827 |
| **MC DS Net** | 1760 | 1415 | 239 | 150 | 56 | 1.59 | 0.901 | 0.804 | 0.85 |
| **DeepFinder** | 1567 | 1362 | 64 | 203 | 20 | 2.22 | 0.867 | 0.869 | 0.868 |
| **DeepETPicker** | 1510 | 1447 | 43 | 118 | 19 | 1.15 | **0.921** | 0.958 | 0.939 |
| **TM-F** | 1772 | 963 | 295 | 601 | 17 | 2.65 | 0.613 | 0.543 | 0.576 |
| **TM** | 4195 | 1073 | 583 | 492 | 716 | 2.62 | 0.683 | 0.256 | 0.372 |
| **TomoSwin3D (ours)** | 1434 | 1420 | 14 | 145 | 6 | 1.47 | 0.907 | **0.99** | **0.947** |

A salient property of TomoSwin3D’s performance is its extremely low false-positive rate. TomoSwin3D produces only 14 false positives, compared to 43 for DeepETPicker and 64 for DeepFinder, while achieving a comparable true positive count (TP = 1420 for TomoSwin3D vs. 1447 for DeepETPicker). This FP reduction drives TomoSwin3D’s near-perfect precision (0.990); the highest among all methods. This indicates that the detections are highly reliable and that the model rarely confuses background structures with protein particles. Such behavior is particularly advantageous in downstream subtomogram averaging (STA) pipelines, where false positives can propagate into 2D/3D classification and degrade reconstruction quality.

In terms of sensitivity, TomoSwin3D achieves the second highest recall of 0.907 with 145 FN, which is slightly lower than DeepETPicker (0.921, FN = 118) but remains among the strongest in the comparison. Importantly, TomoSwin3D’s slight recall trade-off is compensated by the substantial gain in precision, leading to the best harmonic balance (F1 = 0.947). This operating characteristic suggests that TomoSwin3D prioritizes high-confidence particle hypotheses, reducing spurious detections while still recovering the vast majority of true particles.

### 2.3 Performance of TomoSwin3D on SHREC 2020 synthetic test data

On the standard SHREC 2020 test data ^34^ (12 different complexes; weight/shape visualized in **Figure 3A**), TomoSwin3D achieved the highest overall performance, with a mean F1 score of 0.933, outperforming the strongest competing deep learning method DeepETPicker (0.923) and the next-best method UMC (0.905). This corresponds to an absolute improvement of +0.010 over DeepETPicker and +0.028 over UMC in macro-averaged F1, confirming that TomoSwin3D provides the best aggregate detection quality across the full suite of targets. In addition to its top mean score, TomoSwin3D maintained uniformly strong results across all complexes, with F1 values consistently in a narrow high-performance band (0.880 - 0.966), indicating robust behavior across diverse particle morphologies and imaging conditions rather than gains driven by a subset of “easy” cases (**Table 3)**.

**Table 3:** Multiclass particle-detection performance comparison of different methods on the SHREC 2020 (synthetic) test set.

| Methods | 1S3X | 3QM1 | 3GL1 | 3H84 | 2CG9 | 3D2F | 1U6G | 3CF3 | 1BXN | 1QVR | 4CR2 | 4D8Q | Mean F1 |
| --- | --- | --- | --- | --- | --- | --- | --- | --- | --- | --- | --- | --- | --- |
| **3D MS-D** | 0.192 | 0.408 | 0.437 | 0.416 | 0.368 | 0.461 | 0.492 | 0.719 | 0.948 | 0.851 | 0.942 | 0.964 | 0.600 |
| **DeepFinder** | 0.61 | 0.729 | 0.8 | 0.911 | 0.783 | 0.848 | 0.866 | 0.939 | **1** | 0.984 | 0.993 | 0.993 | 0.871 |
| **3D ResNet** | 0.193 | 0.185 | 0.405 | 0.407 | 0.334 | 0.445 | 0.491 | 0.628 | 0.906 | 0.719 | 0.868 | 0.817 | 0.533 |
| **YOPO** | 0.558 | 0.741 | 0.67 | 0.834 | 0.696 | 0.682 | 0.795 | 0.896 | 0.987 | 0.83 | 0.923 | 0.993 | 0.800 |
| **Dn3DUnet** | 0.529 | 0.577 | 0.569 | 0.674 | 0.332 | 0.523 | 0.462 | 0.676 | 0.925 | 0.684 | 0.907 | 0.974 | 0.653 |
| **UMC** | 0.661 | 0.827 | 0.839 | 0.947 | 0.855 | 0.873 | 0.899 | 0.981 | 0.997 | 0.98 | **1** | **0.997** | 0.905 |
| **TM-T** | 0.2 | 0.102 | 0.248 | 0.727 | 0.555 | 0.869 | 0.835 | 0.88 | 0.934 | 0.97 | 0.968 | 0.945 | 0.686 |
| **TM-F** | 0.319 | 0.219 | 0.207 | 0.66 | 0.589 | 0.808 | 0.815 | 0.945 | 0.939 | 0.966 | 0.968 | 0.945 | 0.698 |
| **DeepETPicker** | 0.699 | 0.842 | 0.855 | **0.971** | 0.871 | 0.931 | **0.961** | **0.987** | 0.994 | **0.989** | 0.993 | 0.985 | 0.923 |
| **TomoSwin3D (ours)** | **0.88** | **0.9** | **0.89** | 0.901 | **0.944** | **0.94** | 0.95 | 0.955 | 0.96 | 0.952 | 0.966 | 0.959 | **0.933** |

**Figure 3:**
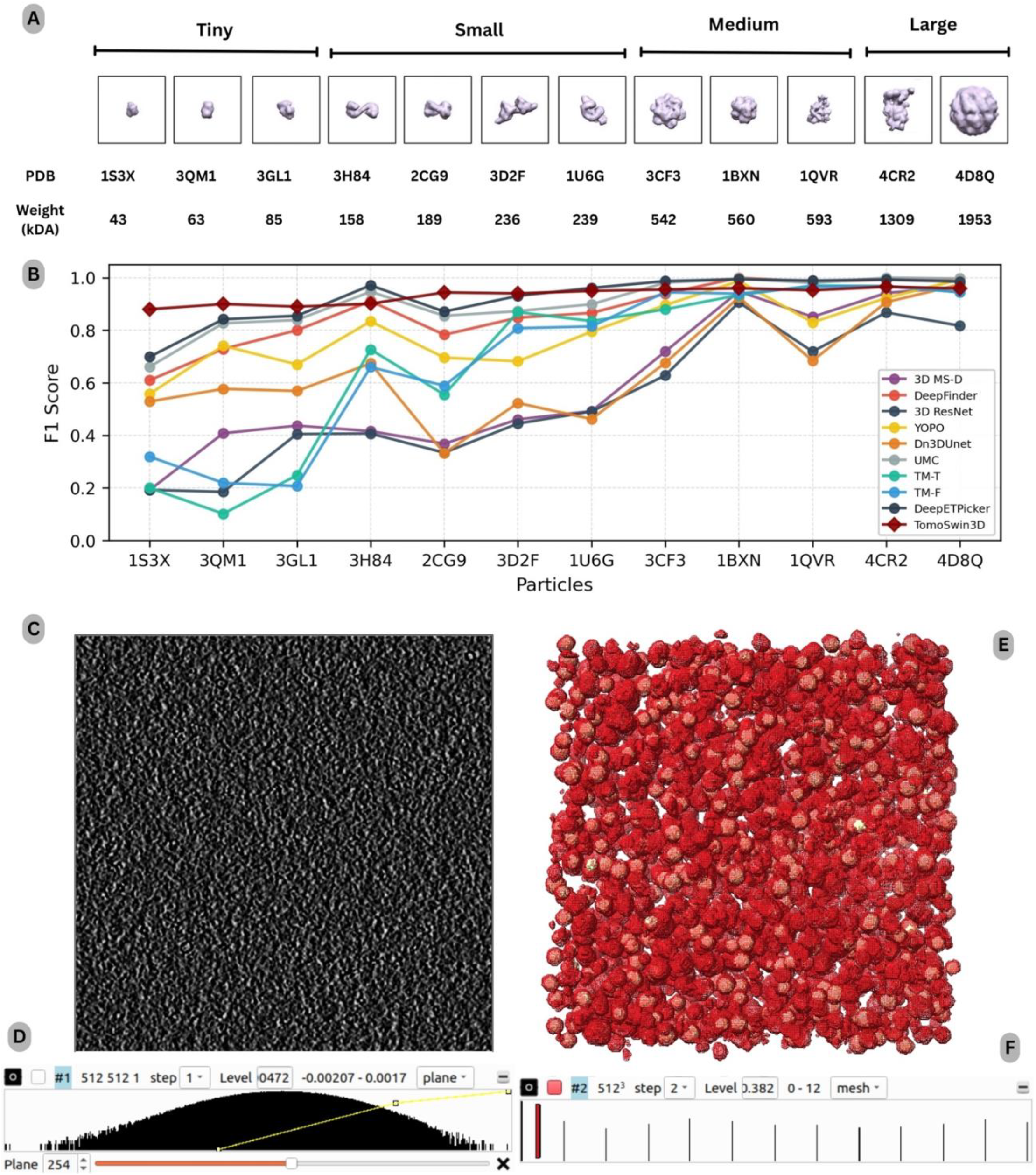
Performance analysis on the SHREC 2020 (synthetic) dataset. **(A)** protein classes in the test set, grouped by size (tiny–large) with corresponding PDB IDs and molecular weights (kDa). **(B)** Per-class F1-score comparison of TomoSwin3D against existing methods for multiclass particle detection across all target proteins. **(C)** Example raw tomogram slice at (z=254), illustrating the low-SNR and texture-dominated background typical of the synthetic reconstructions. **(D)** Voxel-intensity distribution of the input raw tomogram, highlighting the wide dynamic range prior to normalization. **(E)** 3D rendering of the reconstructed volume showing the spatial arrangement with TomoSwin3D predictions (red mesh) overlaid against the reference annotations (golden). **(F)** Volume-viewer rendering (UCSF Chimera) showing the spatial distribution of predicted multiclass macromolecules across the tomogram.

Similarly, as on SHREC 2021 dataset, a key advantage of TomoSwin3D is its pronounced improvement on the most challenging protein targets in this benchmark; those for which several classical and learning-based methods exhibit large performance drops. As shown in Figure 3B, on 1S3X, TomoSwin3D reaches 0.880, exceeding the best competing result (DeepETPicker: 0.699) by +0.181 F1. Similarly, TomoSwin3D delivers the best score on 3QM1 (0.900; +0.058 over DeepETPicker), 3GL1 (0.890; +0.035), and 2CG9 (0.944; +0.073). These cases are typically associated with lower contrast and/or smaller, less distinctive density patterns, and the consistent gains suggest that TomoSwin3D is particularly effective at preserving sensitivity while controlling false positives under low-SNR conditions; precisely where template-based approaches and earlier deep architectures tend to struggle. Detailed metrics are provided in Supplementary Table S5.

For higher-scoring (comparatively easier) protein complexes; where several methods approach saturation - TomoSwin3D remains highly competitive, typically producing F1 ≥ 0.95 (e.g., 1BXN: 0.960, 1QVR: 0.952, 4CR2: 0.966, 4D8Q: 0.959) (see **Supplementary Table S6, S7 & S8** for more macromolecules details). In these cases, minor gaps to the best-performing methods often reflect near-perfect ceiling effects (e.g., UMC at 1.000 on 4cr2 or DeepFinder at 1.000 on 1BXN) rather than substantive detection failures. Overall, the SHREC 2020 results demonstrate that TomoSwin3D provides the best macro-level accuracy while also delivering notable gains on the hardest particle classes, yielding a favorable balance between peak performance and cross-target reliability. As an example, a test tomogram and its raw density values are shown in **Figure 3C&D** and output predictions and multiclass values are shown in **Figure 3E&F**.

To complement the per-protein complex analysis, we further evaluated binary protein particle detection on the SHREC 2020 test set using standard detection metrics (TP/FP/FN, recall, precision, miss rate, F1, and others) as shown in **Table 4**. Across all competing methods, TomoSwin3D achieved the best overall detection accuracy, with an F1 score of 0.9521. This exceeds the strongest baselines UMC (F1 = 0.949) and DeepETPicker (F1 = 0.945), and substantially outperforms earlier deep-learning approaches such as 3D MS-D (0.926), DeepFinder (0.924), and YOPO (0.907), as well as template-matching variants (TM-F: 0.841; TM-T: 0.704) and a generic 3D ResNet baseline (0.702). Overall, these results indicate that TomoSwin3D provides the most favorable balance between sensitivity and specificity on the SHREC 2020 evaluation test dataset.

**Table 4:**
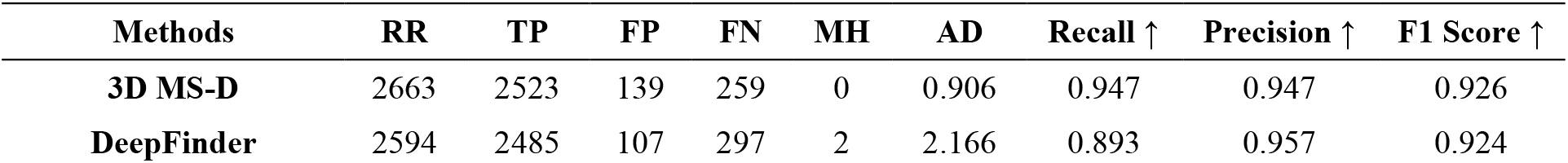
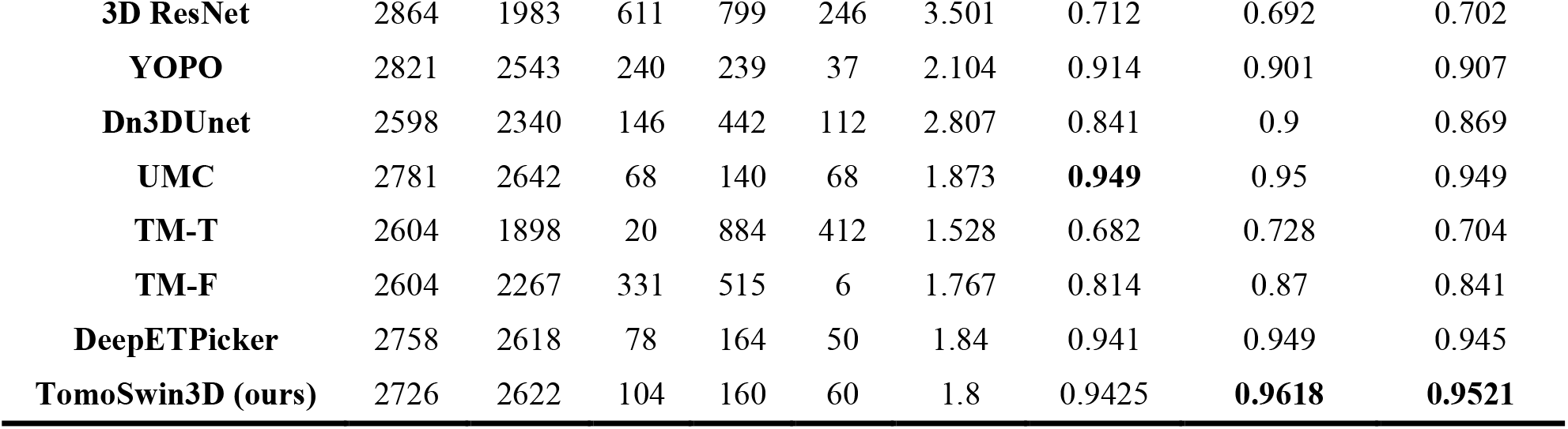
Binary class particle-detection performance comparison of existing methods on the SHREC 2020 (synthetic) test set (RR: reported results i.e., number of particles detected by each method; TP: true positives, representing correctly identified unique particles; FP: false positives, indicating detections of particles that do not exist; FN: false negatives, referring to actual particles that were missed; MH: multiple hits, where a single true particle is detected more than once; AD: mean Euclidean distance (in voxels) between predicted and actual particle centers; Recall: ratio of correctly identified unique particles to the total number of true particles in the test tomogram; Precision: ratio of correctly identified unique particles to the total reported results (RR); F1 Score: harmonic mean of precision and recall).

A closer inspection of the error profile highlights why TomoSwin3D yields the top F1-score. TomoSwin3D attains the highest precision among all methods (0.9618) while preserving high recall (0.9425), resulting in a low miss rate (1-recall = 0.0575). In absolute counts, TomoSwin3D produced TP = 2622, FP = 104, and FN = 160, demonstrating strong control of false positives while maintaining sensitivity. Compared with DeepETPicker, TomoSwin3D improves precision (0.9618 vs. 0.949) and slightly improves recall (0.9425 vs. 0.941), which together translate into a +0.0071 absolute gain in F1-score. Relative to UMC, which attains the highest recall (0.949), TomoSwin3D achieves a higher precision (0.9618 vs. 0.950), yielding a higher overall F1 despite a small reduction in recall. This trade-off suggests that TomoSwin3D is particularly effective at suppressing spurious detections—an important property for downstream subtomogram averaging workflows, where excess false positives can significantly degrade reconstruction quality and increase manual curation burden.

Importantly, TomoSwin3D also exhibits strong localization fidelity, with AD = 1.8, which is among the best values reported for deep learning-based methods in this comparison (e.g., UMC: 1.873; DeepETPicker: 1.84; 3D MS-D: 2.05; DeepFinder: 2.166). Template matching with a tight threshold (TM-T) reports a lower AD (1.528) but suffers from markedly reduced recall (0.682), indicating that improved localization alone is insufficient when sensitivity collapses in low-SNR cellular tomograms. Taken together, the SHREC 2020 binary evaluation demonstrates that TomoSwin3D delivers state-of-the-art detection accuracy while maintaining high-precision predictions and competitive localization, supporting its suitability for reliable, large-scale particle picking in practical cryo-ET pipelines.

### 2.4 Performance of TomoSwin3D on the EMPIAR experimental test data

On the EMPIAR test data (EMPIAR-10731 dataset) consisting of real-world experimental tomograms, TomoSwin3D achieved the strongest overall performance among all compared methods, consistently surpassing CFNPicker, DeepFinder, and DeepETPicker across precision, recall, and F1-score (as shown in **Figure 4A**). For the EMPIAR benchmark, the comparisons were limited to CFNPicker, DeepFinder, and DeepETPicker, as these are among the few methods with publicly available implementations that enable reproducible evaluation on this dataset. In contrast, for the SHREC 2020 and 2021 benchmarks, results from a broader set of methods were included based on previously published reports by SHREC. Since the EMPIAR-10731 dataset contains only a single protein species (ribosomes) across its tomograms, particle detection was formulated as a binary classification task distinguishing particle vs. background voxels. Consequently, multiclass classification was not required for this benchmark, and all compared methods were evaluated under the same binary detection setting. Averaged over the 12 test tomograms (TS_01–TS_12), TomoSwin3D obtained a precision of 0.703, a recall of 0.850, and an F1-score of 0.769. In comparison, the corresponding average F1-scores of CFNPicker, DeepFinder, and DeepETPicker were 0.627, 0.673, and 0.731, respectively. These results indicate that TomoSwin3D provides the most favorable balance between sensitivity and false-positive control on experimentally acquired cryo-ET data.

**Figure 4:**
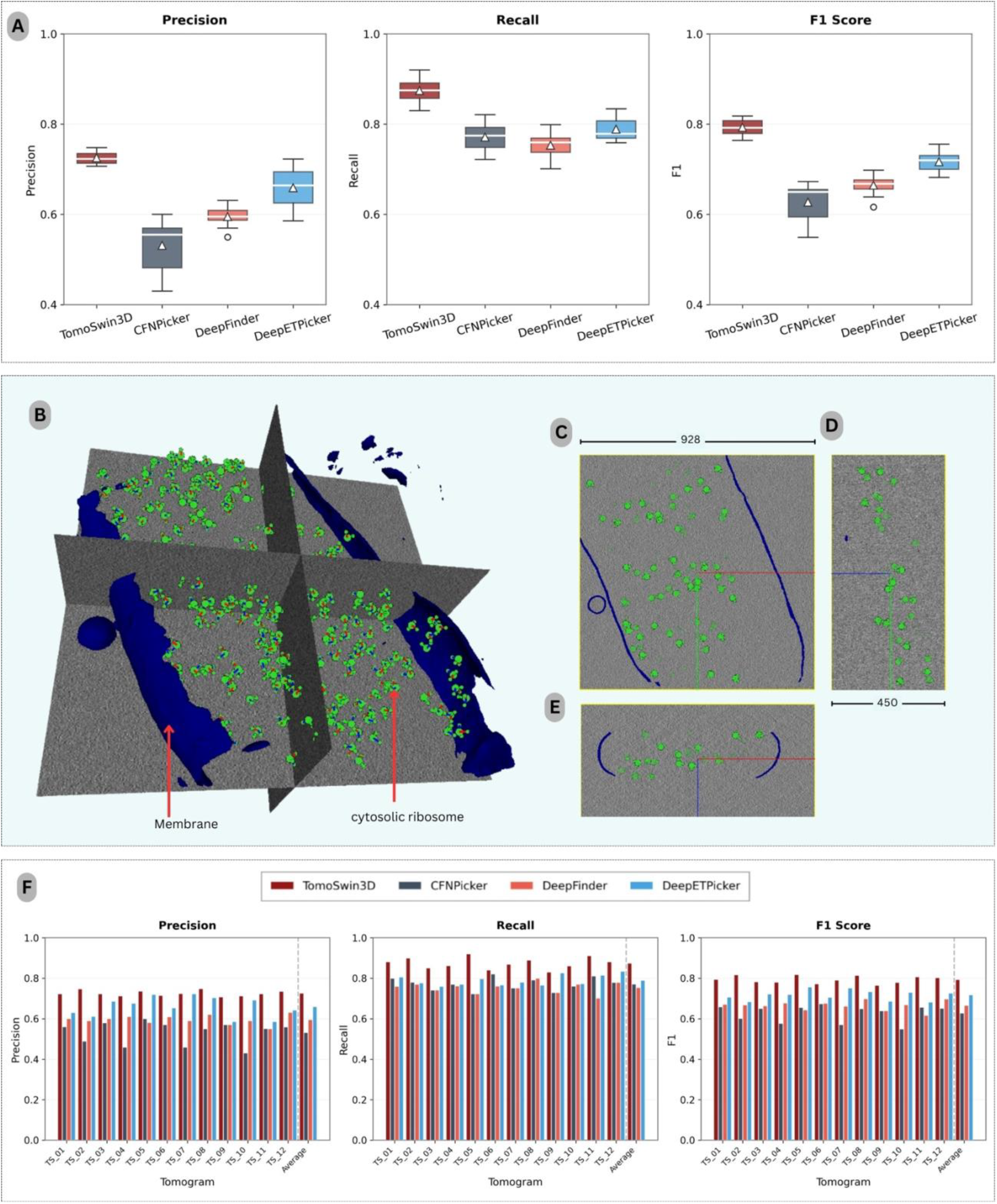
Performance analysis on the experimental EMPIAR-10731 dataset. (A) Box plots summarizing precision, recall, and F1-score across the 12 test tomograms. (B) 3D rendering of a representative input tomogram volume (928 × 928 × 450 voxels), highlighting the segmented membrane (blue) and cytosolic ribosomes (green), shown with orthogonal slice planes for spatial context. (C–E) Orthogonal views of the same volume in the XY, YZ, and XZ planes. (F) Per-tomogram precision, recall, and F1-score bar plots for all 12 test tomograms across methods.

At the individual tomogram level, TomoSwin3D ranked first on 8 of the 12 test tomograms in all three evaluation metrics simultaneously, namely TS_01, TS_02, TS_04, TS_05, TS_06, TS_08, TS_10, and TS_11 (the 3D rendering of a test tomogram and its orthogonal views shown in **Figure 4BCD & E**). Its strongest performance was observed on TS_05, where it achieved a precision of 0.736, a recall of 0.920, and an F1-score of 0.818. Similarly high performance was obtained on TS_02 (F1 = 0.816), TS_08 (F1 = 0.813), and TS_11 (F1 = 0.806). These results suggest that TomoSwin3D is highly effective at recovering true particles while maintaining relatively strong precision, which is critical for reducing the burden of downstream false-positive filtering (the detailed metrics are provided in **Supplementary Table S9, S10, S11 & S12**).

Relative to the competing methods, TomoSwin3D yielded consistent improvements in overall detection accuracy on the EMPIAR dataset. In terms of average F1-score, it exceeded CFNPicker by 0.142 (0.769 vs. 0.627; ∼22.6% relative improvement), DeepFinder by 0.096 (0.769 vs. 0.673; ∼14.3% relative improvement), and DeepETPicker by 0.038 (0.769 vs. 0.731; ∼5.2% relative improvement), as summarized in **Table 5**. (see **Supplementary Table S13** for the true count of particles in each tomogram). Notably, these gains were not obtained through a trade-off between precision and recall. Instead, TomoSwin3D achieved the highest average precision (0.703) and the highest average recall (0.850), outperforming CFNPicker (0.531/0.771), DeepFinder (0.604/0.762), and DeepETPicker (0.672/0.804) in both metrics simultaneously. This pattern indicates that TomoSwin3D more effectively suppresses spurious detections while preserving sensitivity to true particles, thereby producing the most balanced and reliable particle-picking performance across the EMPIAR test dataset.

**Table 5:** Binary particle detection performance comparison of different methods on the EMPIAR-10731 (experimental) test set.

| Methods/Tomogram | TomoSwin3D (ours) |  |  | CFNPicker |  |  | DeepFinder |  |  | DeepETPicker |  |  |
| --- | --- | --- | --- | --- | --- | --- | --- | --- | --- | --- | --- | --- |
|  | Precision | Recall | F1 | Precision | Recall | F1 | Precision | Recall | F1 | Precision | Recall | F1 |
| TS_01 | **0.723** | **0.881** | **0.794** | 0.56 | 0.799 | 0.658 | 0.6 | 0.759 | 0.67 | 0.63 | 0.805 | 0.706 |
| TS_02 | **0.747** | **0.899** | **0.816** | 0.489 | 0.78 | 0.601 | 0.59 | 0.769 | 0.667 | 0.611 | 0.777 | 0.684 |
| TS_03 | 0.686 | 0.759 | 0.721 | 0.579 | 0.741 | 0.65 | 0.6 | 0.741 | 0.663 | **0.723** | **0.85** | **0.782** |
| TS_04 | **0.712** | **0.861** | **0.78** | 0.459 | 0.77 | 0.576 | 0.61 | 0.761 | 0.677 | 0.675 | 0.77 | 0.719 |
| TS_05 | **0.736** | **0.92** | **0.818** | 0.6 | 0.722 | 0.655 | 0.58 | 0.722 | 0.643 | 0.719 | 0.797 | 0.756 |
| TS_06 | **0.714** | **0.84** | **0.772** | 0.57 | 0.821 | 0.673 | 0.609 | 0.76 | 0.676 | 0.653 | 0.766 | 0.705 |
| TS_07 | 0.723 | 0.78 | 0.751 | 0.459 | 0.751 | 0.57 | 0.59 | 0.751 | 0.661 | **0.724** | **0.869** | **0.79** |
| TS_08 | **0.748** | **0.889** | **0.813** | 0.55 | 0.791 | 0.649 | 0.62 | 0.799 | 0.698 | 0.703 | 0.765 | 0.733 |
| TS_09 | 0.586 | 0.826 | 0.686 | 0.57 | 0.729 | 0.639 | 0.57 | 0.729 | 0.639 | **0.707** | **0.83** | **0.764** |
| TS_10 | **0.712** | **0.86** | **0.779** | 0.43 | 0.76 | 0.549 | 0.59 | 0.769 | 0.668 | 0.692 | 0.773 | 0.73 |
| TS_11 | **0.723** | **0.91** | **0.806** | 0.551 | 0.81 | 0.656 | 0.55 | 0.701 | 0.616 | 0.586 | 0.814 | 0.682 |
| TS_12 | 0.631 | 0.779 | 0.697 | 0.559 | 0.779 | 0.651 | **0.735** | **0.881** | **0.802** | 0.642 | 0.834 | 0.726 |
| Average | **0.703** | **0.85** | **0.769** | 0.531 | 0.771 | 0.627 | 0.604 | 0.762 | 0.673 | 0.672 | 0.804 | 0.731 |

Overall, the EMPIAR results demonstrate that TomoSwin3D generalizes robustly to experimental cryo-ET data and delivers the most consistent particle-picking accuracy among the evaluated methods. Its ability to achieve the highest average precision, recall, and F1-score across the tomograms highlights the effectiveness of the proposed framework for real-world experimental cryo-electron tomography.

### 2.5 Performance of TomoSwin3D on the cryo-ET portal test data

Across the three Cryo-ET data portal test tomograms ^35^ (macromolecules shape and size shown in **Figure 5A)**, TomoSwin3D achieved the strongest overall performance among all compared methods whose codebase was openly available for running the predictions. Its average F1-score reached 0.893, exceeding DeepETPicker (0.850), DeepFinder (0.843), and CFNPicker (0.780) (**Figure 5B)**. This improvement was supported by the highest average true positive count (482.33) together with the lowest average false negatives (14.67), indicating that TomoSwin3D recovered more target particles while missing fewer instances than the competing approaches. The method also produced the highest average recall (0.971), demonstrating consistently strong detection sensitivity across all test volumes.

**Figure 5:**
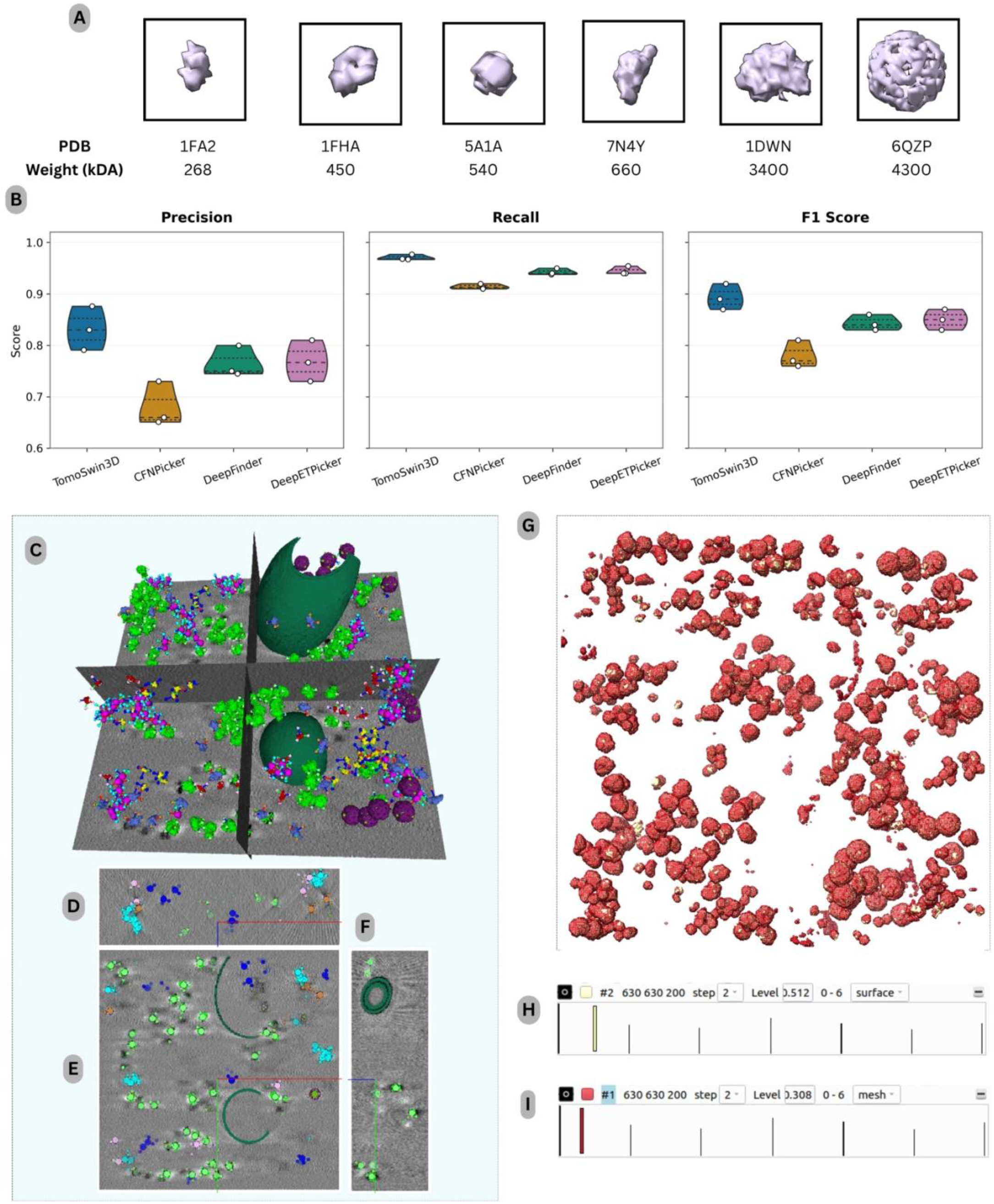
TomoSwin3D performance on Cryo-ET portal test tomograms. (A) Target macromolecules included in the evaluation, shown as representative 3D structures with their PDB identifiers and molecular weights (kDa). (B) Per-method precision, recall, and F1-score for three test tomograms (Tomogram 24-26). (C) 3D rendering of a sample input tomogram with orthogonal slice planes, illustrating the spatial distribution of macromolecules and membranes. (D–F) Orthogonal slice views (XY, XZ, and YZ). (G) Whole-volume 3D visualization of TomoSwin3D predictions (red mesh) overlayed against ground truth particles (golden). (H–I) Volume-viewer rendering (UCSF Chimera) showing the spatial distribution of ground truth particles and predicted multiclass macromolecules across the tomogram.

A tomogram-level analysis (3D rendering of a test tomogram and its orthogonal views shown in **Figure 5CDE & F)** further shows that TomoSwin3D remained the top-performing method on Tomogram_24, Tomogram_25, and Tomogram_26. As shown in **Table 6**, on Tomogram_24, it achieved an F1-score of 0.87, compared with 0.85 for DeepETPicker, 0.83 for DeepFinder, and 0.76 for CFNPicker. On Tomogram_25, its F1-score increased to 0.92, representing the best result and reflecting a favorable balance between high recall (0.968) and strong precision (0.876). On Tomogram_26, TomoSwin3D again led with an F1-score of 0.89 and the highest recall of 0.977, confirming that its advantage was not limited to a single tomogram but was maintained across different test cases (output predictions and multiclass values shown in **Figure 5G, H&I**).

**Table 6:**
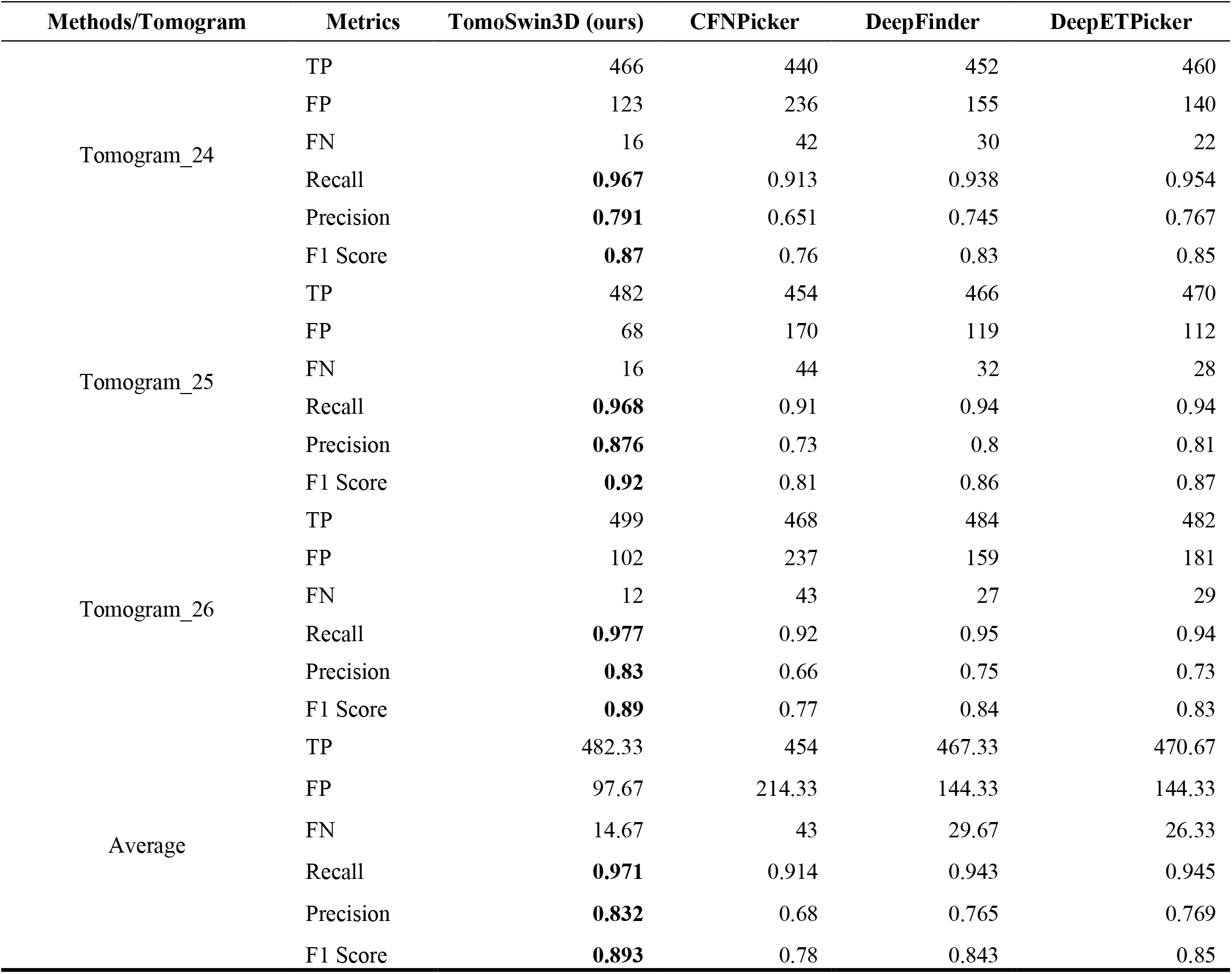
Binary particle detection performance comparison of different methods on the Cryo-ET portal test set (true particle count for tomogram 24, 25 and 26 are: 482, 489, and 511).

| Methods/Tomogram | Metrics | TomoSwin3D (ours) | CFNPicker | DeepFinder | DeepETPicker |
| --- | --- | --- | --- | --- | --- |
| Tomogram_24 | TP | 466 | 440 | 452 | 460 |
|  | FP | 123 | 236 | 155 | 140 |
|  | FN | 16 | 42 | 30 | 22 |
|  | Recall | **0.967** | 0.913 | 0.938 | 0.954 |
|  | Precision | **0.791** | 0.651 | 0.745 | 0.767 |
|  | F1 Score | **0.87** | 0.76 | 0.83 | 0.85 |
| Tomogram_25 | TP | 482 | 454 | 466 | 470 |
|  | FP | 68 | 170 | 119 | 112 |
|  | FN | 16 | 44 | 32 | 28 |
|  | Recall | **0.968** | 0.91 | 0.94 | 0.94 |
|  | Precision | **0.876** | 0.73 | 0.8 | 0.81 |
|  | F1 Score | **0.92** | 0.81 | 0.86 | 0.87 |
| Tomogram_26 | TP | 499 | 468 | 484 | 482 |
|  | FP | 102 | 237 | 159 | 181 |
|  | FN | 12 | 43 | 27 | 29 |
|  | Recall | **0.977** | 0.92 | 0.95 | 0.94 |
|  | Precision | **0.83** | 0.66 | 0.75 | 0.73 |
|  | F1 Score | **0.89** | 0.77 | 0.84 | 0.83 |
| Average | TP | 482.33 | 454 | 467.33 | 470.67 |
|  | FP | 97.67 | 214.33 | 144.33 | 144.33 |
|  | FN | 14.67 | 43 | 29.67 | 26.33 |
|  | Recall | **0.971** | 0.914 | 0.943 | 0.945 |
|  | Precision | **0.832** | 0.68 | 0.765 | 0.769 |
|  | F1 Score | **0.893** | 0.78 | 0.843 | 0.85 |

An important aspect of these results is that the performance gain of TomoSwin3D was driven not only by higher sensitivity but also by better control of false detections. Although DeepETPicker and DeepFinder showed relatively competitive recall values, both methods produced more false positives on average (144.33 each) than TomoSwin3D (97.67), which reduced their precision and overall F1-score. The contrast with CFNPicker was even more pronounced, as its substantially higher false positive rate (214.33) and lower precision (0.680) led to the weakest overall performance. Taken together, these findings indicate that TomoSwin3D provides the most accurate particle picking on the Cryo-ET portal test data.

Because each Cryo-ET portal tomogram contains multiple macromolecular species, a full evaluation would normally involve multiclass detection. However, the existing baseline methods used for comparison were not trained on these specific proteins and therefore cannot assign class identities for these targets. To ensure a fair comparison across methods, we evaluated all methods using binary particle detection as described above. However, for completeness, the multiclass performance of TomoSwin3D on the same dataset is reported separately in **Supplementary Table S14**.

**Supplementary Table S14** summarizes multiclass performance of TomoSwin3D on 3 tomograms from Cryo-ET portal dataset that includes 6 different proteins (PDB IDs: 1DWN, 1FA2, 1FHA, 5A1A, 6QZP, 7N4Y). Across all targets, TomoSwin3D achieved consistently strong detection accuracy, with mean F1 scores of 0.835 (tomogram model_24), 0.871 (tomogram model_25), and 0.864 (tomogram model_26). Averaged across the three tomograms, this corresponds to an overall mean F1 of 0.857, indicating robust performance on Cryo-ET portal data without requiring method-specific tuning per tomogram (see **Supplementary Table S15** for the macromolecule details of Cryo-ET portal data).

The best performance is obtained on pp7_vlp (PDB code: 1DWN, 3400 kDa), with F1 scores tightly clustered between 0.902-0.926 (mean across models 0.912), demonstrating reliable detection for very large, structurally distinctive particles. Similarly, high accuracy is achieved for cytosolic_ribosome (PDB code: 6QZP, 4300 kDa) and beta_galactosidase (PDB code: 5A1A, 540 kDa), both reaching a mean across-model F1 of 0.881 (ranges 0.855-0.899 and 0.852–0.914, respectively). On Thyroglobulin (PDB code: 7N4Y, 660 kDa) strong results were also obtained (mean 0.874, range 0.844-0.909). In contrast, the most challenging target is beta_amylase (1FA2; 268 kDa), on which the consistently lowest F1 across all checkpoints (0.738-0.803, mean 0.774) were produced, followed by ferritin_complex (1FHA; 450 kDa) (mean 0.816, range 0.800-0.840). Notably, 1FA2 remains the lowest-scoring class for every checkpoint, suggesting that small proteins with reduced spatial footprint and weaker contrast in crowded cellular environments remain intrinsically harder to localize.

## 3. Discussion

Across four independent benchmarks, TomoSwin3D delivers a strong and reliable 3D particle detection performance across both simulated and experimental cryo-ET data, with its most important advantage being not only higher average F1-score but also better results on the hardest protein targets. On both SHREC datasets, TomoSwin3D achieved the best overall macro-level accuracy, reaching 0.861 on SHREC 2021 and 0.933 on SHREC 2020, outperforming prior deep-learning methods such as DeepETPicker, MC DS Net, CFNPicker, DeepFinder, and UMC. The gains are especially meaningful because they are observed in benchmarks where several competing methods already perform strongly on easy classes; thus, the improvement is not simply due to ceiling effects on easy targets but rather reflects better detection for hard targets under more difficult conditions.

These targets are likely harder because of weaker contrast, smaller or less distinctive structural signatures, and greater ambiguity from local cellular context. The gains on such protein particles suggest that TomoSwin3D is more effective at capturing discriminative multi-scale 3D contextual information, allowing it to maintain sensitivity while limiting false positives.

The results on the real-world experimental tomograms in the EMPIAR-10731 test dataset further affirm the generalizability of TomoSwin3D. On EMPIAR, TomoSwin3D outperformed all other methods in average precision, recall, and F1-score simultaneously, and achieved the best F1-score on 8 out of 12 test tomograms. Its lower cross-tomogram variability compared with prior methods indicates improved robustness to changing imaging conditions, specimen composition, and experimental noise. On Cryo-ET portal, TomoSwin3D also performed strong across 6 biologically diverse protein targets and three independent tomograms, with an overall mean F1 of 0.857.

Taken together, these findings indicate that the main strength of TomoSwin3D lies in its combination of high average accuracy, robustness across varying tomograms and protein shape/sizes, and improved performance on intrinsically difficult detection scenarios. At the same time, the Cryo-ET portal results reveal an important remaining limitation: small proteins such as 1FA2 remain consistently harder to detect than larger and more distinctive complexes, likely due to weaker contrast, and stronger susceptibility to missing-wedge and low-SNR effects. This suggests that, although TomoSwin3D advances automated cryo-ET particle picking, further improvement may depend on strategies specifically tailored to very small-particle localization, such as enhanced small-object supervision. Overall, the present results support TomoSwin3D as a robust and generalizable framework for cryo-ET particle detection, with clear advantages for downstream sub-tomogram extraction and structural analysis workflows.

Despite the transformative impact of deep learning on particle picking in cryo-ET tomograms in recent years, several fundamental challenges persist that constrain the throughput and accuracy of cryo-ET-based visual proteomics. The primary limitation of supervised architectures remains the annotation burden. Voxel-wise segmentation models typically require thousands of high-quality manual labels to train, a process that is prohibitively labor-intensive and subjective. While weakly supervised strategies and Positive-Unlabeled (PU) learning have reduced this requirement to a fraction of the total dataset, these methods often rely on the accurate estimation of class prior probabilities, where minor inaccuracies can lead to significant increases in false-positive or false-negative rates. More advanced deep learning methods require much less labeled training data such as self-supervised learning, zero-shot learning, or few-shot learning may help mitigate the problem.

As revealed in this study, the detection of small macromolecules (<200 kDa) remains a formidable “small object” problem to a certain extent, which is also the case in other 3D computer vision fields. At these scales, protein signals are frequently overwhelmed by extremely low signal-to-noise ratios (SNR) and irregular background noise, such as reconstruction artifacts from weighted back-projections. Many current architectures struggle to preserve weak structural features as they are attenuated through deep convolutional layers. Furthermore, standard grid-cell-based detection often fails to distinguish between adjoining particles in crowded cellular environments due to resolution limits in the grid. More sensitive deep learning architectures and more data containing small protein particles are needed to further address this challenge.

Domain generalization presents another critical hurdle. Models trained on synthetic datasets often exhibit performance degradation when applied to different cell types or novel imaging conditions. This “synthetic-to-real” gap is exacerbated by variability in missing-wedge artifacts, ice thickness, and defocus gradients. Additionally, while current models excel at identifying globular complexes, most lack the specialized mechanisms required to pick filamentous assemblies or membrane-bound proteins without extensive geometric pre-processing.

Future research may prioritize the development of 3D foundation models and universal pickers that utilize zero-shot inference or advanced promptable segmentation to identify novel proteins without per-species retraining. Bridging the data gap will require leveraging large-scale standardized resources to train models on a more diverse range of experimental artifacts. Future particle picking pipelines would also benefit from incorporating denoising algorithms directly into the detection framework to streamline the transition to subtomogram averaging. Rotation-equivariant designs and 3D instance segmentation that explicitly separate aggregated particles are also promising directions for crowded cellular environments and missing-wedge–driven anisotropy.

## 4. Methods

### 4.1 Data processing and feature extraction

#### 4.1.1 Data sources and dataset statistics

We compiled multi-source cryo-ET data comprising both simulated and experimental tomograms to train and test our methods. The simulated dataset includes SHREC 2021 ^31^ and SHREC 2020 ^34^ (both from the “Classification in Cryo-Electron Tomograms” track from SHREC competition), Cryo-ET Data portal DS-10441^35^, and an annotated tomogram collection from the Serpico project team (referred to as Serpico dataset) ^15^. To assess generalization under real cellular imaging conditions, we further evaluated the trained model on experimental EMPIAR-10731 ^36^ dataset as an independent blind test.

Across datasets, each tomogram is available as a reconstructed density volume (MRC format) together with ground-truth annotations. Ground truth is primarily available as multiclass masks (semantic labels), and in addition, particle centroid coordinates (*x, y, z*) are provided to support coordinate-based supervision and downstream evaluation.

The SHREC 2021 dataset contains 10 tomograms (9 for training, 1 for testing), each with volume size 512 × 512 × 512 at 1Å voxel spacing. SHREC 2020 follows the same split (9 training, 1 testing) and volume size (512^3^) at 1 Å voxel spacing. Cryo-ET portal DS-10441 dataset contains 27 tomograms (3 held out for testing), with volumes of 630 × 630 × 200 at 10Å voxel spacing. The Serpico dataset contains 88 tomograms, with volumes of 512 × 512 × 200 at 10.2Å voxel spacing and was used for training purpose. For EMPIAR-10731 dataset, the archive provides tilt series from Mycoplasma pneumoniae cells (12 tilt series) acquired using SerialEM, reported with a tilt range of −60^°^ to +60^°^ in 3^°^ increments and a cumulative exposure of 129e^−^/Å^2^, following an initial pre-exposure of 32 e^−^/Å^2^. The dataset also includes a protein particle segmentation volume and corresponding particle center coordinates. The EMPIAR-10731 dataset was exclusively used for independent testing and was not used for training the model.

In total, we used 126 tomograms for training and 21 tomograms for testing, which includes 12 experimental tomograms from EMPIAR-10731 as an external test set.

#### 4.1.2 Tomogram resampling, intensity normalization, and coordinate alignment

Because the raw tomograms originate from multiple sources with heterogeneous voxel spacings, we standardized spatial resolution by resampling all density volumes to a uniform voxel size of 1Å using the *vol resample* command in UCSF ChimeraX ^37^ (non-interactive mode). The resampled tomograms were then used for subsequent normalization and training.

To mitigate cross-tomogram intensity variability, we applied percentile-based scaling followed by clipping. For a tomogram volume *V*, we compute the 95th percentile of its informative density values, denoted *p*_95_(*V*), and normalize as:

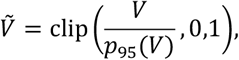

where clip(⋅,0,1) truncates values below 0 to 0 and values above 1 to 1 . This strategy reduces the influence of extreme outliers while enforcing a consistent dynamic range across datasets, thereby facilitating learning of features that are stable across simulation and experimental reconstruction pipelines.

We additionally harmonized coordinate conventions to ensure centroid annotations align with the reconstruction volumes used for training. In the SHREC datasets, the reconstructed tomogram has dimensions 512 × 512 × 512, whereas the ground-truth “*grandmodel*” spans 512 × 512 × 200 and is centered within the full reconstruction. Consequently, an offset must be applied to reconcile the *z*-axis coordinate systems. Following the organizer-provided convention, slice 156 in the reconstruction corresponds to slice 0 in the grandmodel; we therefore applied dataset-specific z-offset correction for SHREC 2020 (z=z+156) and SHREC 2021 (z=z+166) as specified by the SHREC annotations. No offset correction was required for the Cryo-ET portal and Serpico datasets. All coordinates were stored as floating-point values.

#### 4.1.3 Mask generation and label construction

##### 4.1.3.1 Multiclass mask generation

Different sources provide different class ID conventions. We therefore built a tomogram-specific mapping table that remaps each dataset’s original class IDs into a unified multiclass ID space used by our training pipeline. Proteins that appear in some datasets but not others were merged rather than dropped, to preserve supervision and reduce fragmentation in the label space. In addition, vesicles and membranes were excluded from multiclass training because they are not macromolecule particles and would confound a protein-centric detection objective. We implemented multiclass standardization by rewriting each tomogram’s *class_mask* into a new standardized class mask standardized across all datasets, where voxel values are replaced according to the mapping table while preserving the original voxel spacing and origin metadata.

Instead of generating spherical or Gaussian blobs around the particle centroid, our occupancy representation directly uses the original 3D segmentation shape (occupancy) from the class mask - e.g., all voxels belonging to a protein instance retain the mapped class ID {*0,…,K*} (0 = background, K = protein ID), forming a contiguous labeled region (“lump”). This avoids introducing hand-tuned radii or shape assumptions and preserves the original 3D segmented shape of each protein instance.

##### 4.1.3.2 Binary mask generation

In this formulation, we collapse all protein particle labels into a single foreground class: protein voxels are set to 1, and background voxels remain 0, producing a {*0,1*} mask. This binary label construction follows the same occupancy (“lump voxels”) principle described above, and similarly excludes vesicle/membrane regions when present.

### 4.4 Subtomogram (grid) generation

To enable scalable training on full tomograms, we split each normalized 3D tomograms into overlapping cubic subtomogram (“grids”). We use a core grid size of 48×48×48 voxels and add 8-voxel padding on each face to preserve local context ^38^, resulting in a window size of 64×64×64 voxels per cube.

To handle boundaries, the tomogram is padded with zeros prior to splitting so that each grid has a consistent shape. Each grid is saved as a compressed .npz file containing the grid tensor and reconstruction metadata (grid origin indices *i,j,k*, original shape, voxel size, and MRC header orientation fields), which allows reproducible alignment between tomogram cubes and corresponding label cubes.

Tomograms are highly sparse: most 3D subtomogram contain only background noise. To reduce wasted computation and prevent our model from being dominated by background-only samples, we optionally filter training subvolumes using a non-zero grid strategy. Specifically, after splitting a mask into fixed-size cubes, we retain only those cubes whose mask contains at least one non-zero voxel (i.e., grid.max() > 0). This removes completely empty regions of input tomogram while keeping all protein-containing regions for training.

#### 4.1.5 Tomogram feature extraction and multi-channel input

To improve particle detectability under the low SNR and strong background heterogeneity typical of cryo-ET, we augmented each normalized tomogram with three 3D feature maps that emphasize (i) blob-like particle responses across scales, (ii) particle boundaries/edges, and (iii) local contrast enhancement by removing slow intensity trends. All feature volumes were computed directly in 3D on the normalized tomogram and while preserving voxel size metadata and the original origin information. The generation of the three feature maps is described below.

##### 4.1.5.1 Difference-of-Gaussians (DoG) blob response (multi-scale)

We computed a multi-scale DoG response to highlight approximately spherical/ellipsoidal particle-like structures over a range of unknown particle radii. For each scale *s* ∈ {1,2,4,8} voxels, two Gaussian-smoothed volumes were generated using isotropic standard deviations *s* and *ks*, where 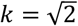. The DoG response was defined as:

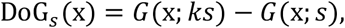

followed by optional scale normalization by *s*^2^ (LoG-approximation scale normalization) to balance responses across particle sizes. At each voxel, we selected the DoG value from the scale that yielded the maximum absolute magnitude, producing a single “blobness” feature volume for protein particles:

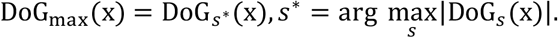

Finally, to stabilize numeric range and reduce sensitivity to outliers, we clipped the resulting map to the [0.01, 99.99] percentile interval.

##### 4.1.5.2 3D Sobel gradients and gradient magnitude

To explicitly encode boundary and edge information, we computed 3D Sobel derivatives along the *x, y*, and *z* axes using nearest-neighbor boundary handling to avoid padding artifacts. Let *g*_*x*_, *g*_*y*_, *g*_*z*_ denote the Sobel gradients. Because voxel spacing can be anisotropic, we converted gradient units from “intensity per voxel” to “intensity per Å” by dividing each component by the corresponding voxel size (*s*_*x*_, *s*_*y*_, *s*_*z*_) read from the MRC header: 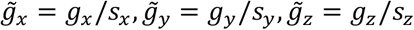.

We then computed the gradient magnitude:

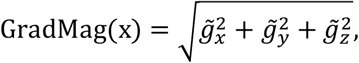

which serves as a rotationally less sensitive edge-strength descriptor.

##### 4.1.5.3 Multi-scale morphological top-hat enhancement

To enhance local particle contrast while suppressing gradual background variations, we computed both white and black top-hat transforms over multiple structuring element sizes. For each scale *s* ∈ {1,2,4,8} voxels, a 3D structuring element (default setting: “ball”-type neighborhood) was used to compute: White top-hat: *T*_*w*_ = *I* − (*I* ° *B*_*s*_), emphasizing bright objects on a darker background (common cryo-ET contrast scenario) and Black top-hat: *T*_*b*_ = (*I* · *B*_*s*_) − *I*, emphasizing dark objects on brighter background.

To gently harmonize responses across scales, each response was multiplied by *s*^1/2^. Across scales, we aggregated responses using a voxel-wise maximum (default setting), producing: (i) a max white top-hat map, (ii) a max black top-hat map, and (iii) a combined top-hat map defined as the maximum of the two polarities at each voxel (and then maximized across scales). As in the DoG pipeline, we clipped each aggregated map to the [0.01,99.99] percentile interval to reduce extreme values.

For the training, each tomogram subvolume was represented as a four-channel tensor by stacking the normalized intensity volume with the three derived feature maps:

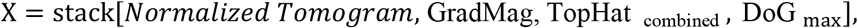

yielding an input of shape (4,64,64,64) per training sample. This multi-view representation supplies the model with important cues-raw density, edge strength, local contrast enhancement, and multiscale blobness-thereby improving robustness to contrast variability and background noise in cryo-ET tomograms.

### 4.2 TomoSwin3D architecture

TomoSwin3D is a 3D Swin Transformer integrated into a U-Net-style encoder-decoder for voxel-wise protein particle classification in cryo-ET (**Figure 6**). The architecture is designed to (i) learn hierarchical representations through downsampling, (ii) model contextual dependencies using localized 3D self-attention within windows, and (iii) recover fine spatial detail through upsampling and multi-scale fusion via skip connections. The overall design maintains computational tractability for volumetric inputs and enabling strong context aggregation, which is critical for particle identification under low SNR and crowded cellular environments.

**Figure 6:**
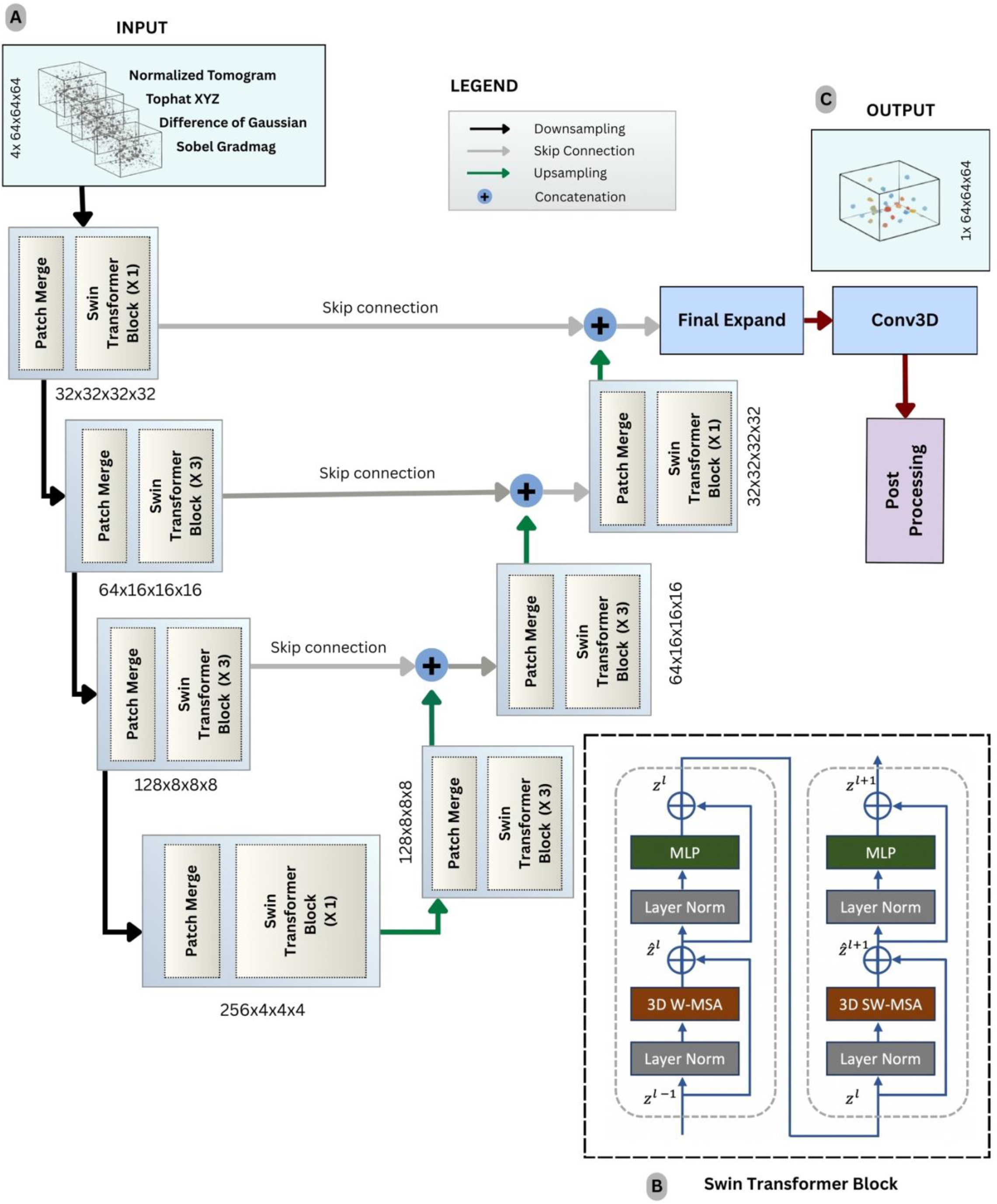
Architecture of TomoSwin3D. **(A)** Input subtomograms (grids) of size 64×64×64 fed into the first encoder block. **(left blocks)** Each encoder block comprises a patch merge layer that downsamples the spatial dimensions and increases the channel dimensions, followed by a Swin Transformer block that includes a window attention mechanism and a multi-layer perceptron (MLP). **(right blocks)** Each decoder block contains a patch-expanding layer that upsamples the input volume and reduces the channel dimensions, followed by another Swin Transformer block. **(B)** TomoSwin3D includes three encoder blocks, a bottleneck block, and three decoder blocks. Output from the decoders is concatenated with corresponding encoder outputs through skip connections for feature propagation. **(C)** The final output is generated via a 3D convolutional block. Predicted outputs grids are stitched back together and post-processed. Detailed Swin Transformer block is illustrated on the bottom right, consisting of 3D window multi-head self-attention (3D W-MSA) block and a 3D shifted window multi-head self-attention (3D SW-MSA) block, each accompanied by an MLP block.

#### 4.2.1 Model overview and input-output specification

Let an input subvolume be denoted by 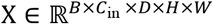, where *B* is batch size, *C*_in_ is the number of input channels, and (*D, H, W*) are the 3D spatial dimensions. In our implementation, each sample corresponds to a 64 × 64 × 64 grid (grid_size = 64) with four input channels (*C*_in_ = 4), obtained by stacking the normalized tomogram intensity and three feature volumes (Sobel gradient magnitude, combined top-hat enhancement, and DoGbased blob response) as shown in **Figure 6A**. The network produces dense per-voxel class logits Z ∈ ℝ^*B*×*K*×*D*×*H*×*W*^, where *K* is the number of semantic classes (e.g., multi-class particle labels plus background). A final 1 × 1 × 1 convolution maps decoder features to Z, and voxel-wise probabilities are obtained through softmax over the class dimension.

Architecturally, TomoSwin3D follows a four-stage encoder and a four-stage decoder, followed by a final upsampling head to restore the original resolution. Each stage combines (1) a resolution transformation module (PatchMerging3D for downsampling or PatchExpand3D for upsampling) and (2) a stack of Swin Transformer blocks that alternate regular and shifted window attention to enable cross-window information flow.

#### 4.2.2 Encoder design

##### 4.2.2.1 PatchMerging3D: learnable volumetric downsampling

Downsampling is implemented using PatchMerging3D, which reduces resolution by a factor *s* (default *s* = 2) along each axis. For an input feature tensor F ∈ ℝ^*B*×*C*×*D*×*H*×*W*^, the volume is partitioned into non-overlapping *s* × *s* × *s* blocks. Each block is flattened into a token of length *C* ⋅ *s*^3^ and projected to a higher-dimensional embedding space via a linear layer, followed by normalization:

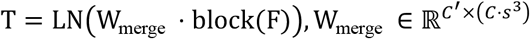

The merged tokens are then reshaped back into a lower-resolution 3D grid for subsequent transformer processing. This operation is analogous to strided convolution in CNNs but preserves a token representation suitable for attention.

##### 4.2.2.2 Multi-stage hierarchical feature extraction

After patch merging, each encoder stage applies a stack of SwinBlock3D units, enabling hierarchical abstraction as depth increases (details of swin transformer block shown in **Figure 6B**). Spatial resolution decreases while channel capacity increases, producing coarse but semantically rich representations at deeper stages. In the default configuration, downscaling is performed with factors (2,2,2,2) across stages, and the base embedding width is hidden_dimension = 32.

Because the architecture alternates regular and shifted attention blocks, each stage depth is typically chosen to be even. A representative depth schedule is (2,6,6,2) across encoder stages, with a corresponding increase in the number of attention heads (3,6,12,24), and a per-head dimensionality head_dimension = 32, supporting high-capacity attention at lower resolutions.

#### 4.2.3 3D Swin transformer blocks

##### 4.2.3.1 Window-based multi-head self-attention in 3d

Standard global self-attention scales quadratically with the number of tokens, which is prohibitive for 3D grids. TomoSwin3D adopts window-based attention, restricting self-attention to local 3D windows of a window_size (default (2,2,2)). Internally, features are converted to channel-last format for window partitioning and attention computation, i.e., ℝ^*B*×*D*×*H*×*W*×*C*^. For each window, queries, keys, and values are computed using a single linear projection:

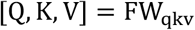

Multi-head attention within a window is then computed as:

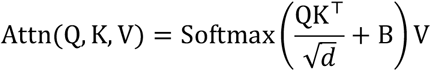

where *d* is the per-head dimension and B is a learnable relative position bias that encodes voxel-pair offsets inside the 3D window.

##### 4.2.3.2 Shifted windows for cross-window interaction

Window attention is efficient but local. To enable interactions across neighboring windows, TomoSwin3D alternates standard window attention with shifted window attention. A cyclic shift by *δ* = window_size /2 is applied along each axis before partitioning; attention is computed within these shifted windows; then the shift is reversed. Since shifting introduces artificial wrap-around adjacency, an attention mask is applied to suppress invalid token interactions across boundaries. This mechanism provides cross-window information flow while preserving the favorable complexity of local attention.

##### 4.2.3.3 Relative position encoding in 3D

To maintain spatial awareness without absolute positional embeddings, TomoSwin3D uses relative positional encoding. A learnable bias table is indexed by discrete 3D offsets within a window and added to the attention logits. This is especially beneficial in cryo-ET volumes, where spatial relations among voxels are crucial for distinguishing particle patterns from background structures.

##### 4.2.3.4 Block structure and feed-forward network

Each SwinBlock3D follows a pre-normalized residual design: 1) LayerNorm → WindowAttention3D → residual add and 2) LayerNorm → MLP (FeedForward3D) → residual add. The MLP expands the channel dimension by a factor of 4 with GELU activation:

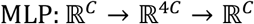

Dropout is applied in the feed-forward path to regularize training, which is often necessary for volumetric transformer models trained under limited annotated data.

#### 4.2.4 Decoder design and skip fusion

##### 4.2.4.1 PatchExpand3D: learnable upsampling

The decoder mirrors the encoder using PatchExpand3D modules. After each upsampling step, SwinBlock3D stacks refine the features at the new resolution, enabling context-aware reconstruction of voxel-level predictions.

##### 4.2.4.2 Multi-Scale skip connections and fusion strategy

To preserve high-frequency spatial detail and stabilize gradient flow, TomoSwin3D uses U-Net skip connections between corresponding encoder and decoder stages. Let F_enc_ and F_dec_ denote encoder and decoder features at the same resolution. Skip fusion is implemented through a configurable module (*skip_style*), with additive fusion as the default:

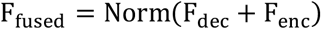

#### 4.2.5 Final upsampling and prediction head

After the final decoder stage, the network applies a final expansion module (FinalExpand3D) to restore the original grid resolution. This step typically includes linear projection, spatial rearrangement, normalization, and a nonlinearity. A concluding 1 × 1 × 1 convolution block (**Figure 6C**) maps these features to class logits:

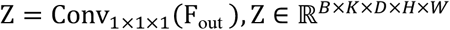

#### 4.2.6 Training objective and optimization details

##### 4.2.6.1 Voxel-wise cross-entropy loss function

Training is performed with voxel-wise supervision using cross-entropy (CE) as the primary objective. For one-hot ground truth labels **y**_*v,k*_ and predicted probabilities *p*_*v,k*_ at voxel *v*, the CE loss is:

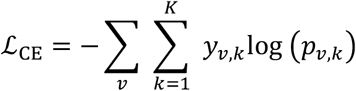

Class weights are applied to address the severe imbalance between background and particle voxels. For our implementation, weights are derived from voxel frequency statistics (e.g., per-batch class counts), giving higher weights to rarer particle classes.

##### 4.2.6.2 Optimization stability: gradient clipping and multi-gpu training

Optimization is carried out using Adam with configuration-driven hyperparameters. For stability in volumetric transformer training, global-norm gradient clipping is applied:

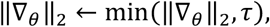

where *τ* is the clipping threshold. The implementation supports multiGPU execution through data parallelism, and checkpoints are saved after each epoch, including model state, optimizer state, training curves, best validation performance, and a snapshot of the full configuration to ensure reproducibility.

### 4.3 Inference and post-processing

#### 4.3.1 Patch-wise inference on test tomograms and reconstruction of full-volume prediction masks

Inference is performed on a test set which inputs 3D sub-volumes (grids) extracted from each tomogram. For each test tomogram, the inference script loads all grids and iterates through them using the model weights.

Given an input grid x, the network outputs per-voxel logits z(v) ∈ ℝ^*C*^ at each voxel position v, where *C* denotes the number of classes (binary or multi-class, depending on the trained checkpoint). For multiclass inference, posterior probabilities are computed using softmax:

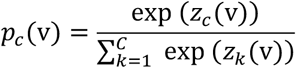

The predicted class label is obtained via 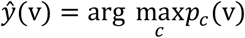. To suppress uncertain predictions, a global confidence threshold *τ* is applied to the maximum posterior:

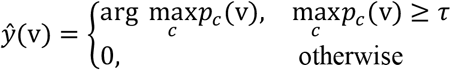

where class 0 denotes background. For binary inference, the particle probability *p*_1_(v) is thresholded to produce a binary mask. Each predicted grid is saved as an .npz file together with its spatial placement metadata (*i, j, k*), effective extents (*d*_*i*_, *d*_*j*_, *d*_*k*_), padding size, and MRC header fields (voxel size, origin, axis mapping).

Because inference is performed patch-wise, a full tomogram-sized prediction volume is reconstructed by stitching the predicted grids back into the original tomogram coordinate frame. For each grid, only its central, non-padded region is used during reconstruction to minimize boundary artifacts. Concretely, if the grid contains a padding margin of *p* voxels, then the reconstructed sub-volume corresponds to grid [*p* : *p* + *d*_*i*_, *p*: *p* + *d*_*j*_, *p*: *p* + *d*_*k*_], which is placed into the global volume at [*i*: *i* + *d*_*i*_, *j*: *j* + *d*_*j*_, *k*:*k* + *d*_*k*_] . The reconstructed prediction is finally written as an MRC volume, while preserving voxel size, origin, and axis mapping information from the original tomogram metadata to maintain downstream spatial consistency.

#### 4.3.2 Morphological post-processing, coordinate extraction and evaluation

The reconstructed output MRC contains discrete voxel labels (background and one or more protein classes). Post-processing converts this dense semantic volume into a sparse list of particle coordinates through three steps: connected-component extraction, size-based filtering, and centroid computation. When ground-truth particle locations are available, predicted centroids are compared to reference coordinates using one-to-one assignment with the Hungarian algorithm under a distance cutoff (5 voxels). Let 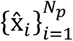 be predicted coordinates and 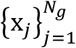 ground truth coordinates. A cost matrix of Euclidean distances 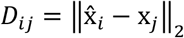 is computed and distances greater than a threshold *d*_max_ are invalidated (set to a large constant), after which linear assignment is solved to obtain an optimal one-to-one matching. Matched pairs within *d*_max_ are counted as true positives (TP), with FP = *N*_*p*_ − TP, FN = *N*_*g*_ − TP. We finally reported standard detection metrics: Precision, Recall, and F1 score for each of the test dataset for performance comparison with existing methods.

## Conclusion

In this work, we developed TomoSwin3D, an end-to-end 3D protein particle picking pipeline that integrates standardized tomogram preprocessing, occupancy-preserving supervision (using instance masks instead of heuristic Gaussian labels), and a Swin3D U-Net that leverages long-range context for dense prediction, followed by simple connected-component post-processing for centroid extraction. Across synthetic and real-world benchmarks (SHREC 2020/2021, EMPIAR, and Cryo-ET portal), TomoSwin3D achieves strong and consistent detection accuracy, demonstrating both competitive performance and improved robustness across heterogeneous tomograms.

TomoSwin3D can reduce the bottleneck in cryo-ET particle picking and support higher-throughput subtomogram analysis, enabling more scalable visual proteomics workflows. Remaining challenges include reliable detection of extremely small/low-contrast macromolecules in crowded cellular environments.

## Data availability

The SHREC 2021 and SHREC 2020 dataset are available on the website of the SHREC 2021 challenge (https://www.shrec.net/cryo-et/) and 2020 challenge (https://www.shrec.net/cryo-et/2020/) respectively. Cryo-ET Data portal DS-10441^35^ can be accessed at https://Cryo-ETdataportal.czscience.com/datasets/10441. The tomogram and annotation from the experimental dataset of Mycoplasma pneumoniae cells can be obtained under accession number EMPIAR-10731 (ref. ^36^). The dataset from Serpico project team (referred to as Serpico dataset) can be accessed through ref. ^15^.

## Code availability

The scripts, programs, and instructions for downloading and installing TomoSwin3D are found in https://github.com/jianlin-cheng/TomoSwin3D.

## Author contributions

J.C. conceived and supervised this research. A.D. and J.C. designed the method. A.D. collected and processed data, implemented, trained, and tested the method. A.D., R.G., and J.C. analyzed the data and results. A.D. and R.G. drafted the manuscript. A.D., R.G., and J.C. edited and revised the manuscript.

## Supplementary information

**Supplementary Table S1:**
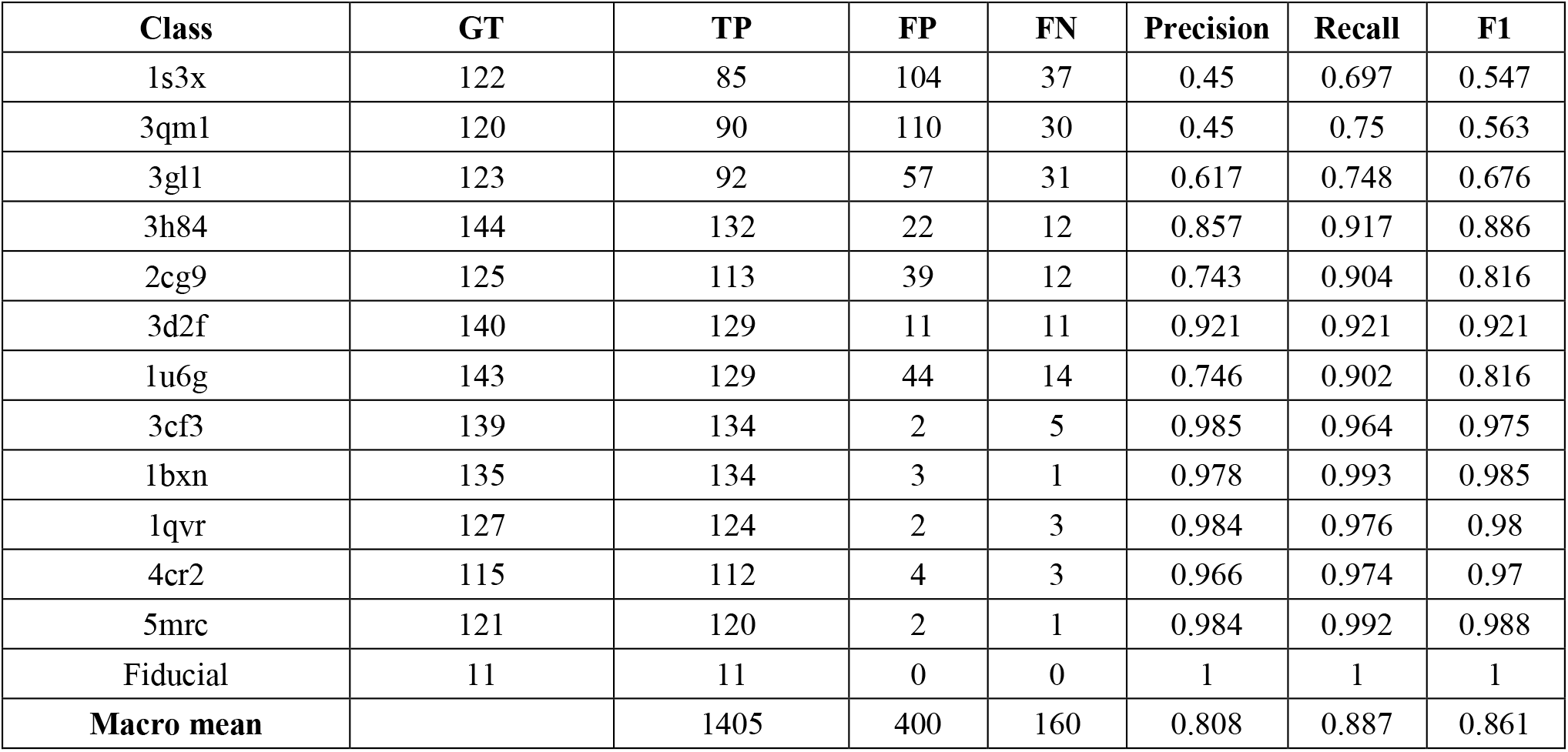
Multiclass particle detection performance metrics of TomoSwin3D on the SHREC 2021 (synthetic) test data.

**Supplementary Table S2:**
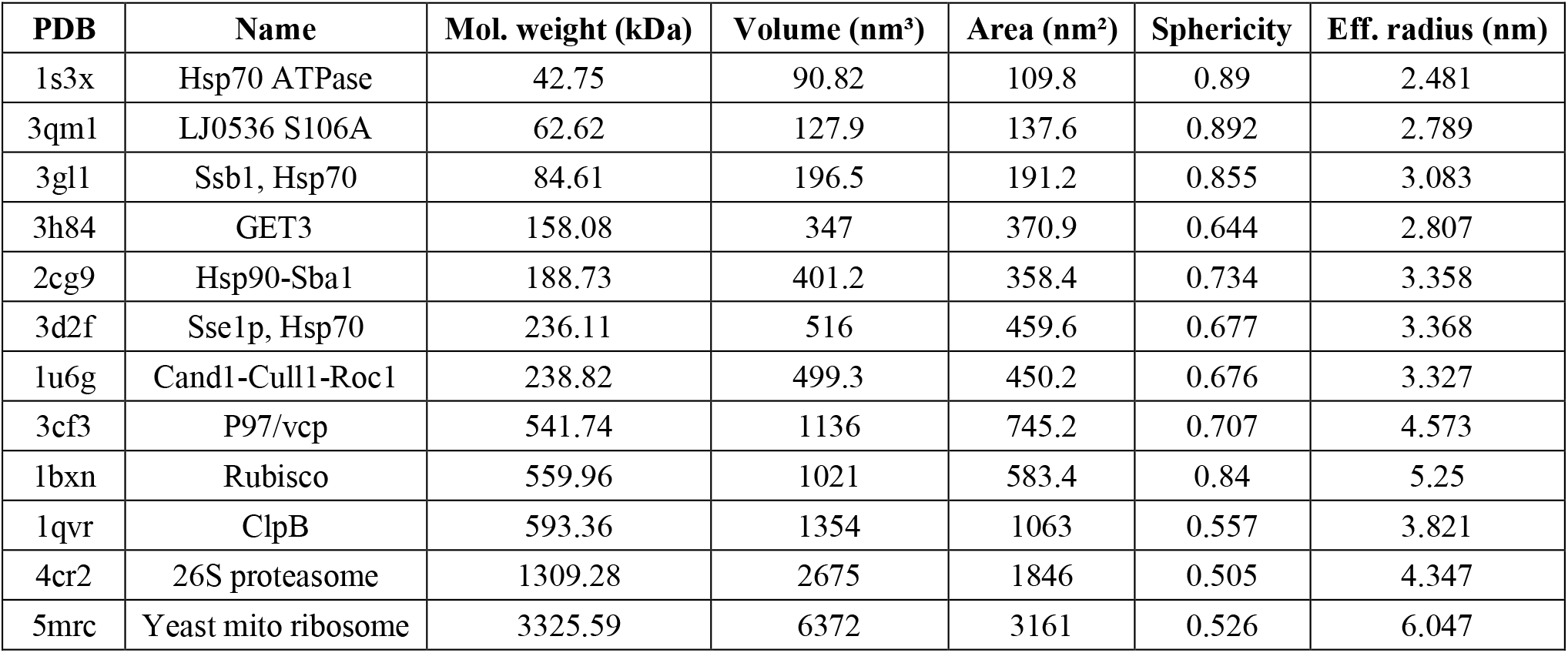
Details of macromolecular complexes present in the SHREC 2021 dataset, sourced from SHREC 2021 competition.

**Supplementary Table S3:**
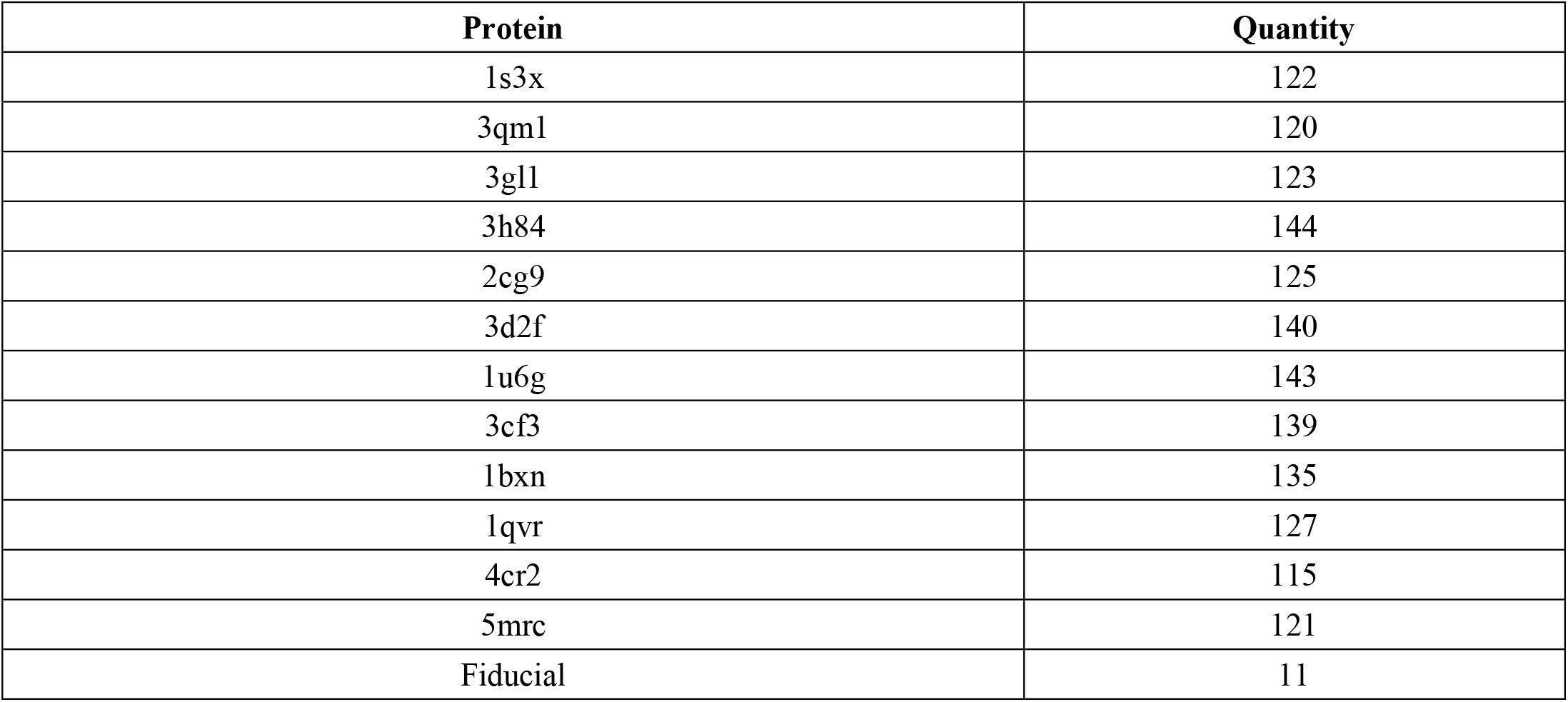
Protein IDs and their numbers present in the SHREC 2021 dataset, sourced from SHREC 2021 competition.

**Supplementary Table S4:**
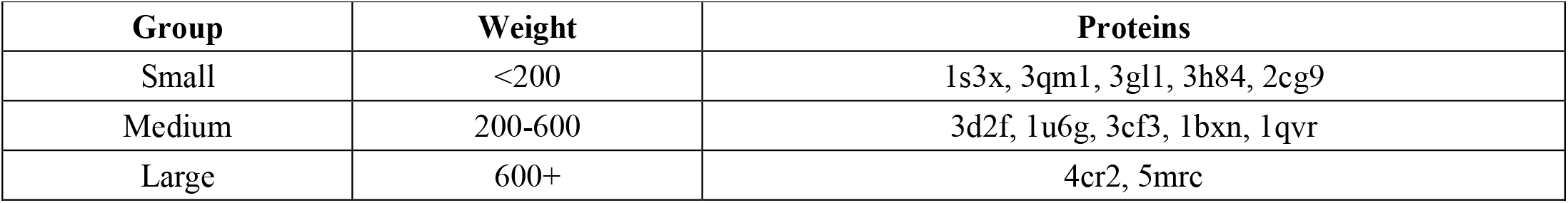
Macromolecular complexes by their molecular weight in kDa, sourced from SHREC 2021 competition.

**Supplementary Table S5:**
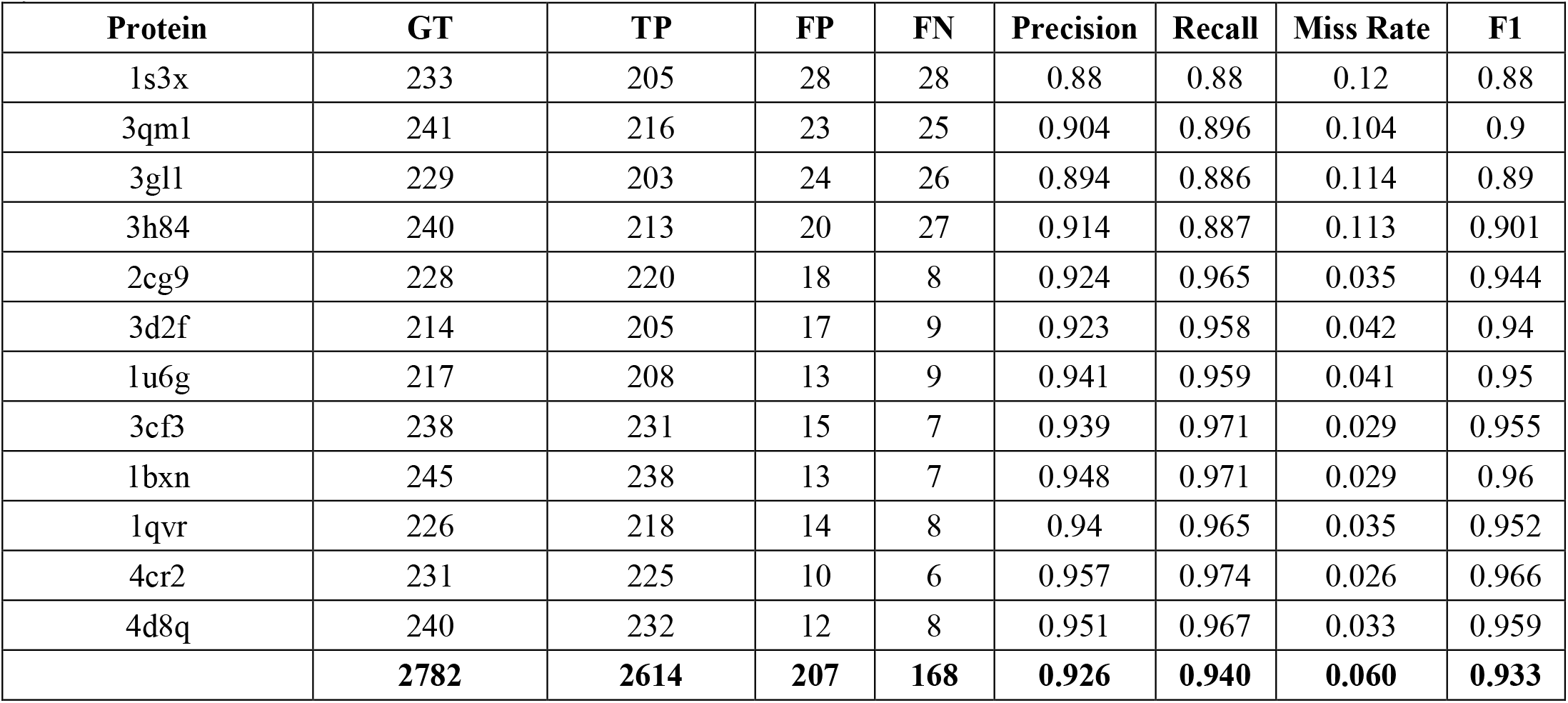
Multiclass particle detection performance metrics of TomoSwin3D on the SHREC 2020 (synthetic) test data.

**Supplementary Table S6:**
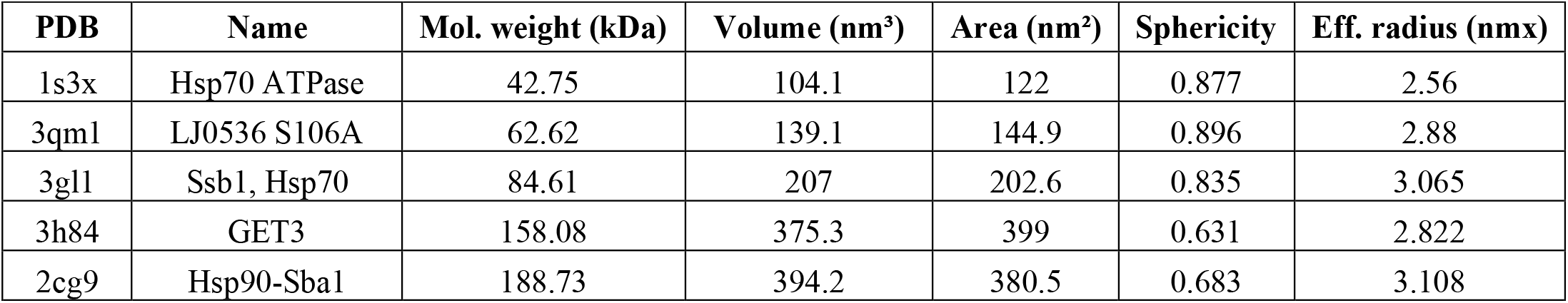
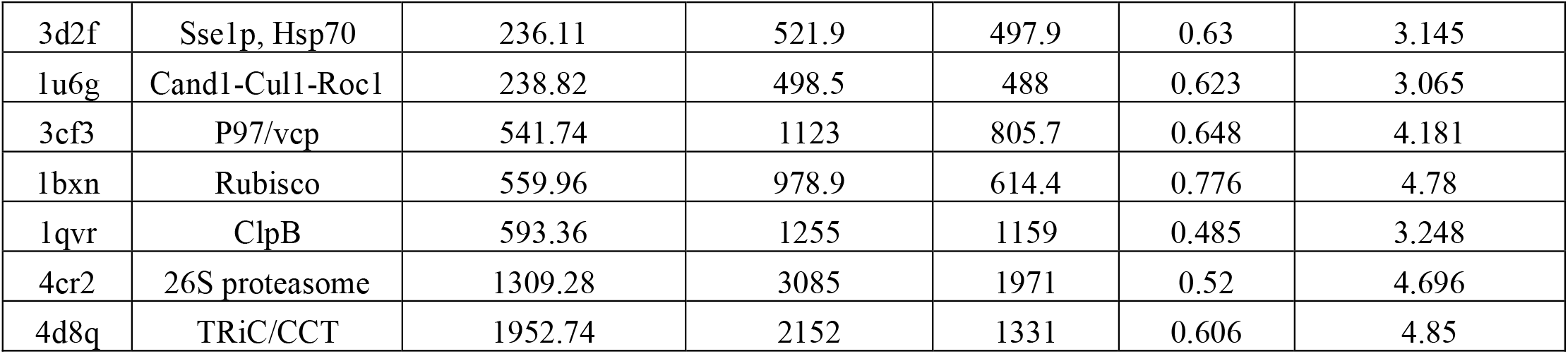
Details of macromolecular complexes present in the SHREC 2020 dataset, sourced from SHREC 2020 competition.

**Supplementary Table S7:**
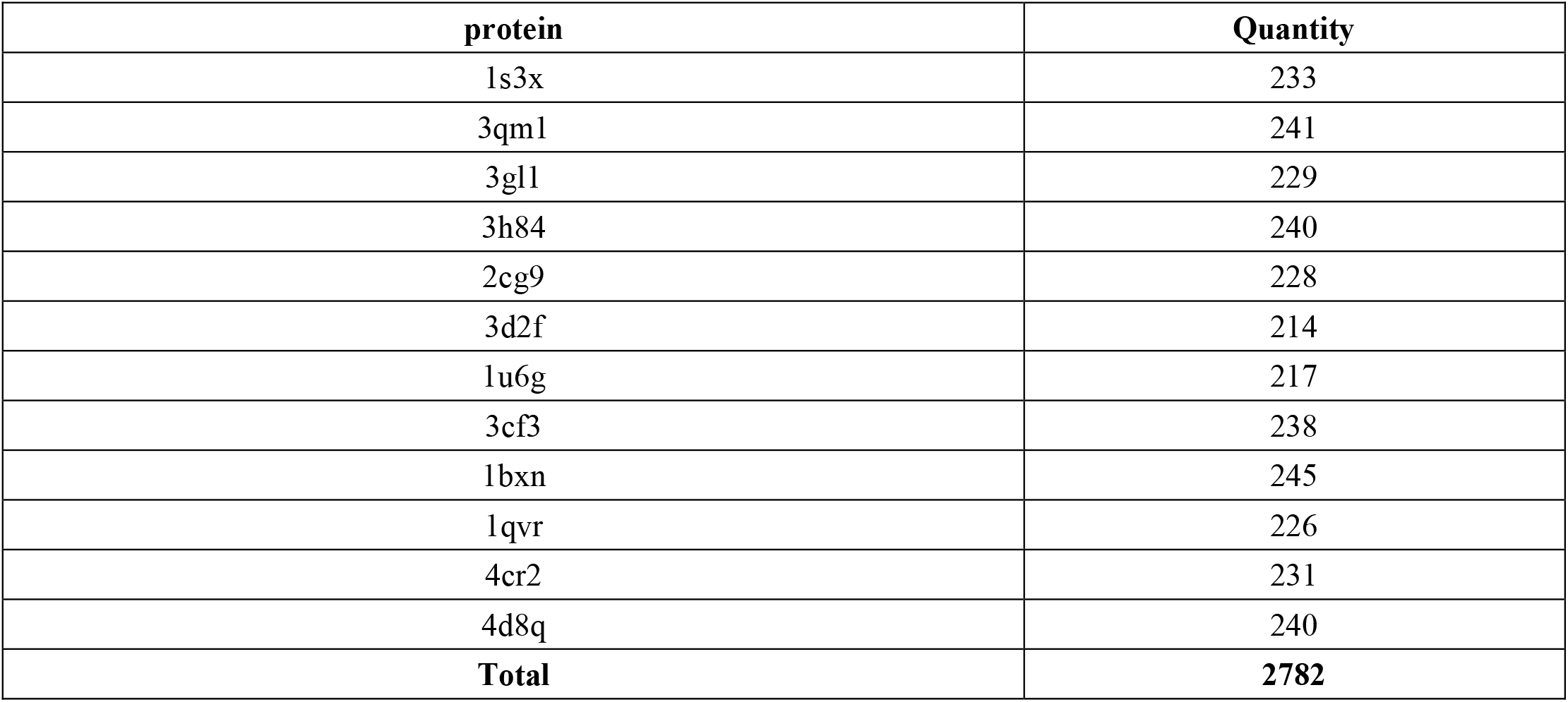
Protein IDs and their numbers present in the SHREC 2020 dataset, sourced from SHREC 2020 competition.

**Supplementary Table S8:**
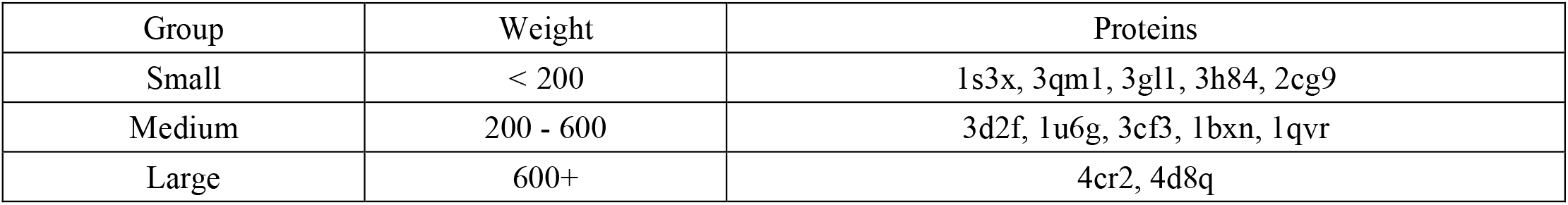
Macromolecular complexes by their molecular weight in kDa, sourced from SHREC 2020 competition.

**Supplementary Table S9:**
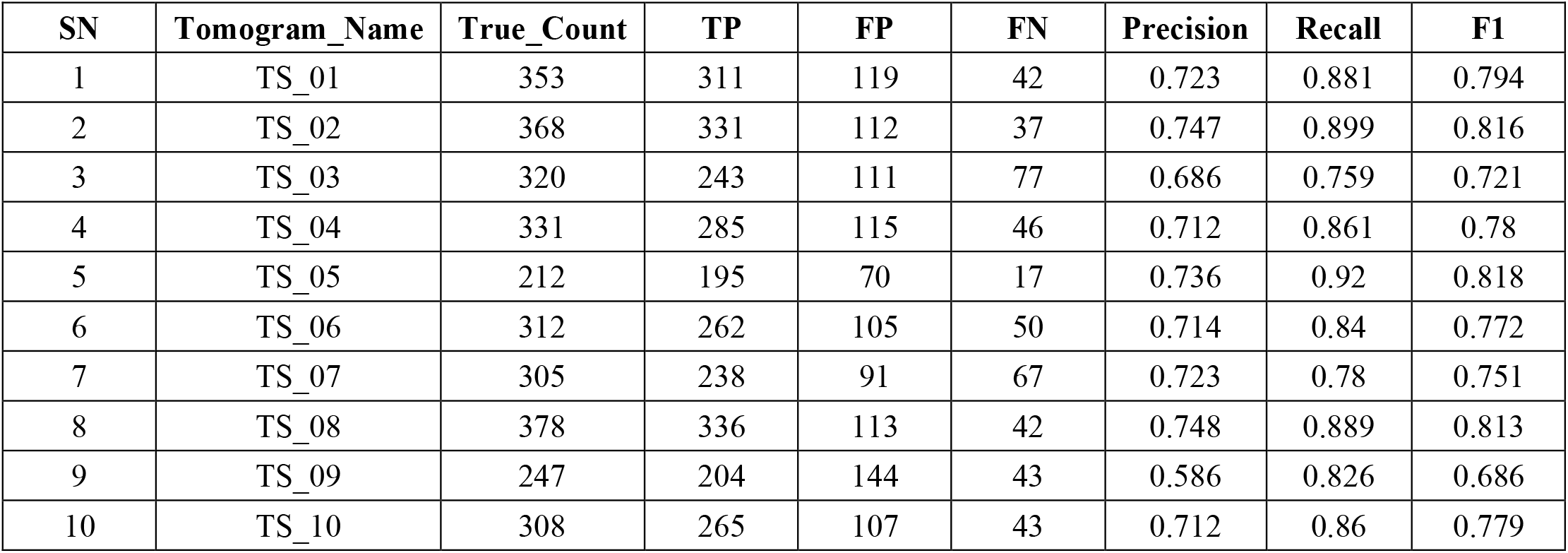
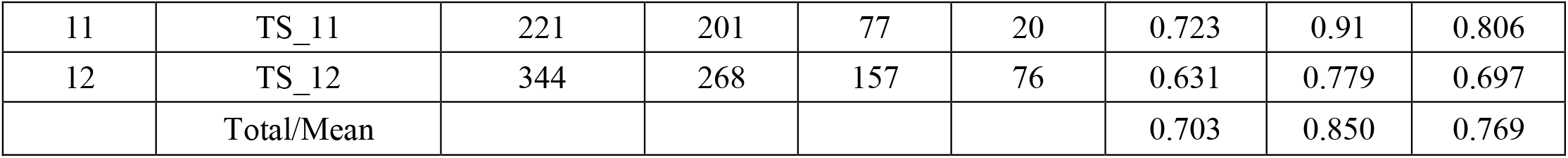
Binary particle detection performance metrics of TomoSwin3D on the EMPIAR-10731 (experimental) test data.

**Supplementary Table S10:**
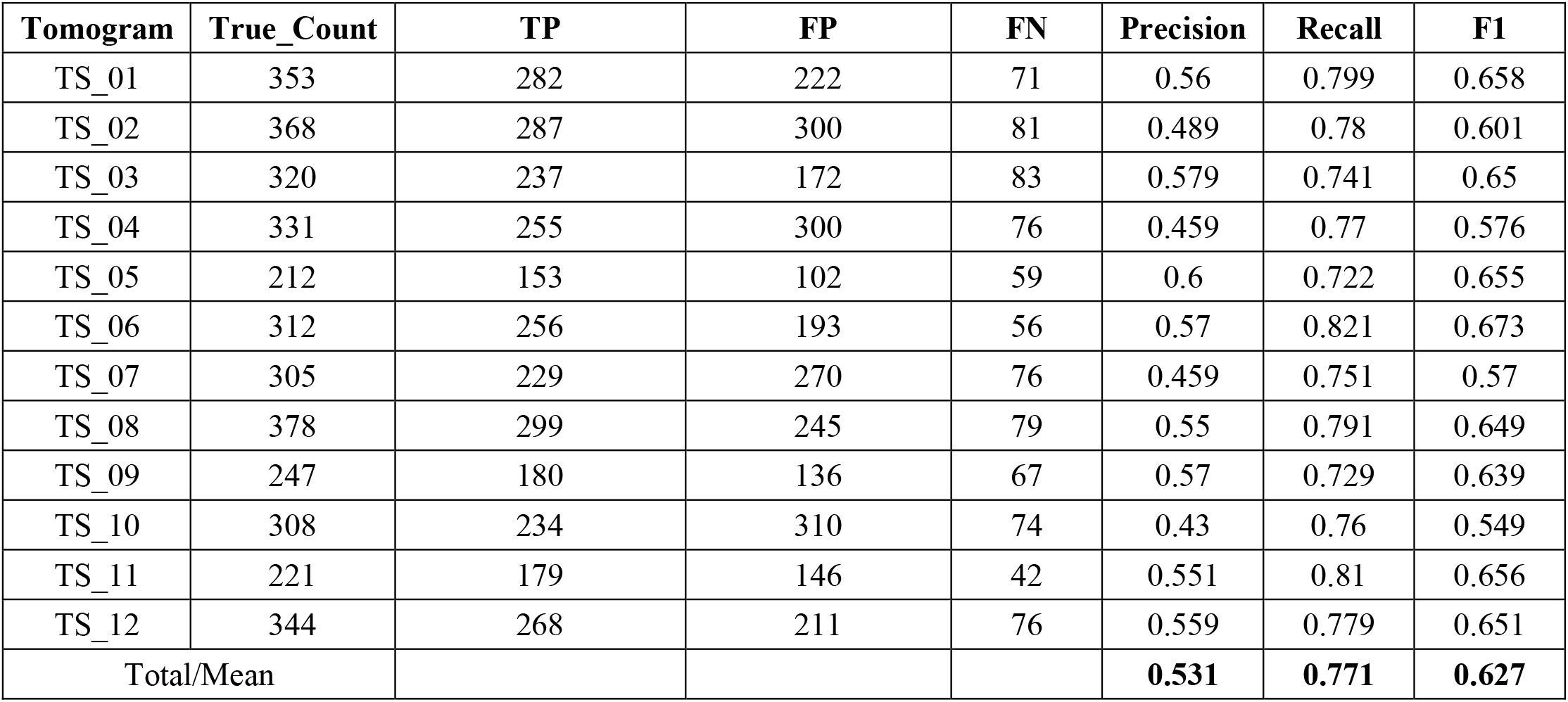
Binary particle detection performance metrics of CFNPicker on the EMPIAR-10731 (experimental) test data.

**Supplementary Table S11:**
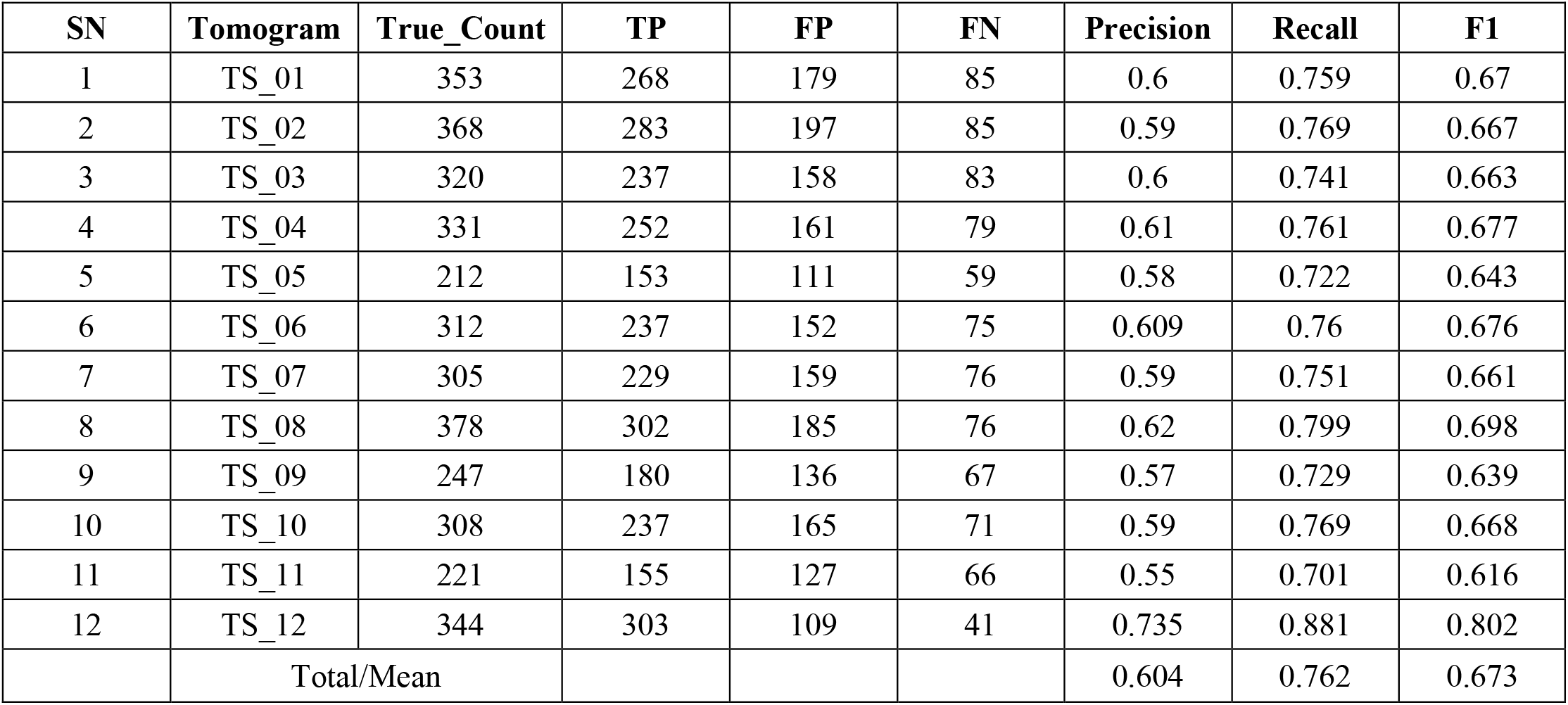
Binary particle detection performance metrics of DeepFinder on the EMPIAR-10731 (experimental) test data.

**Supplementary Table S12:**
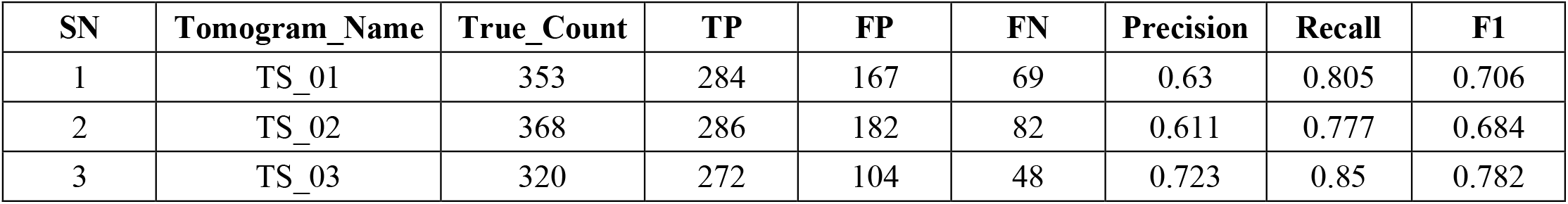
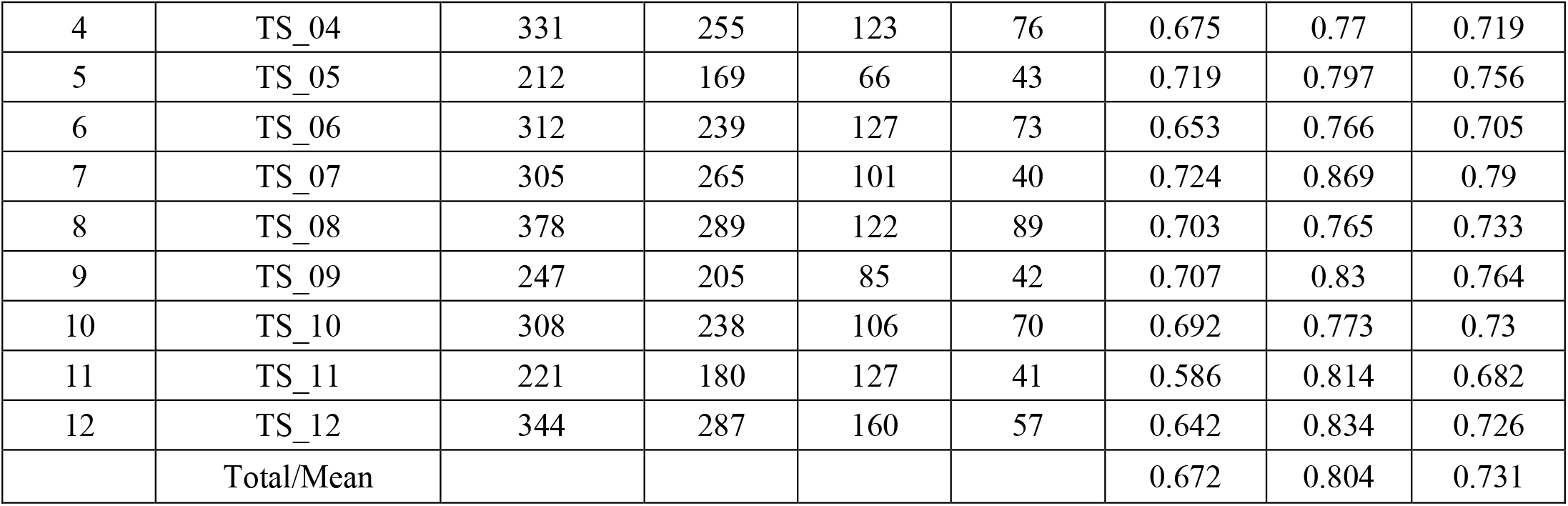
Binary particle detection performance metrics of DeepETPicker on the EMPIAR-10731 (experimental) test data.

**Supplementary Table S13:**
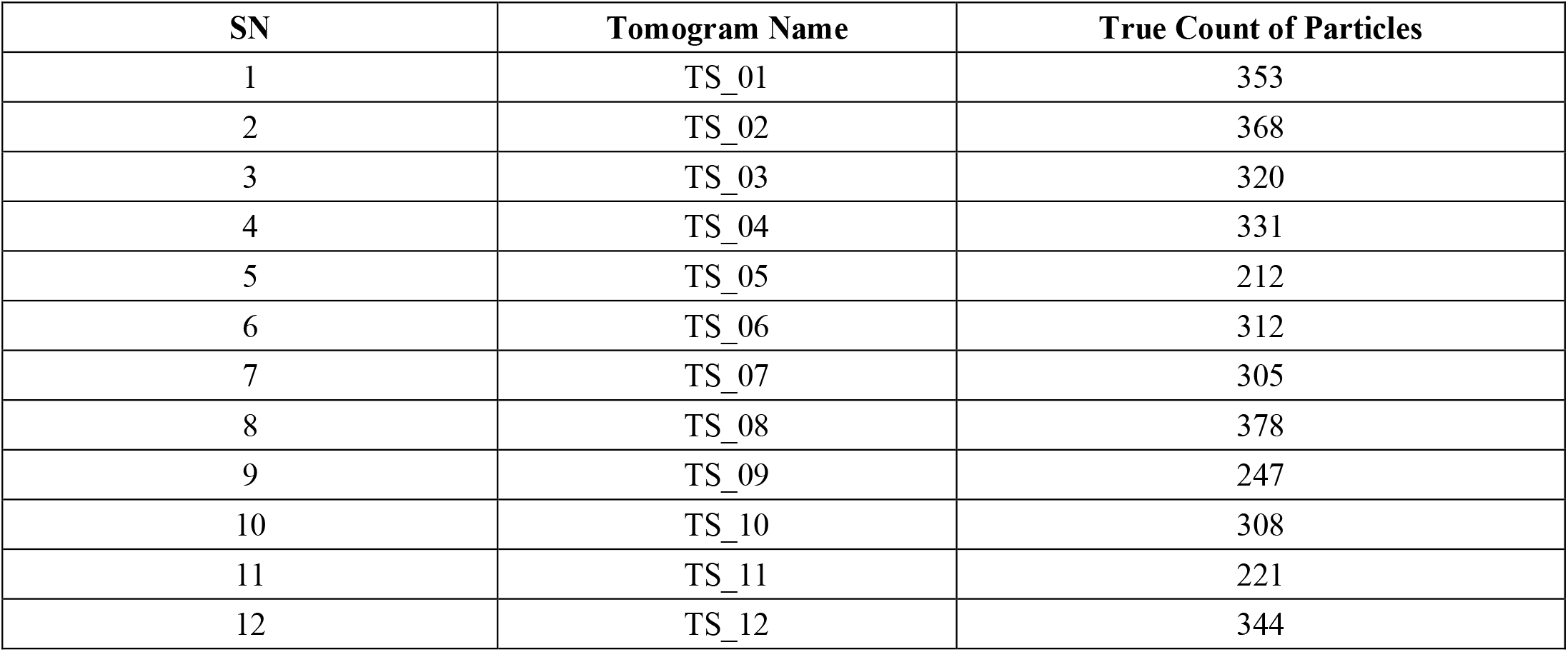
Protein IDs and their numbers present in the EMPIAR-10731 test dataset.

**Supplementary Table S14:**
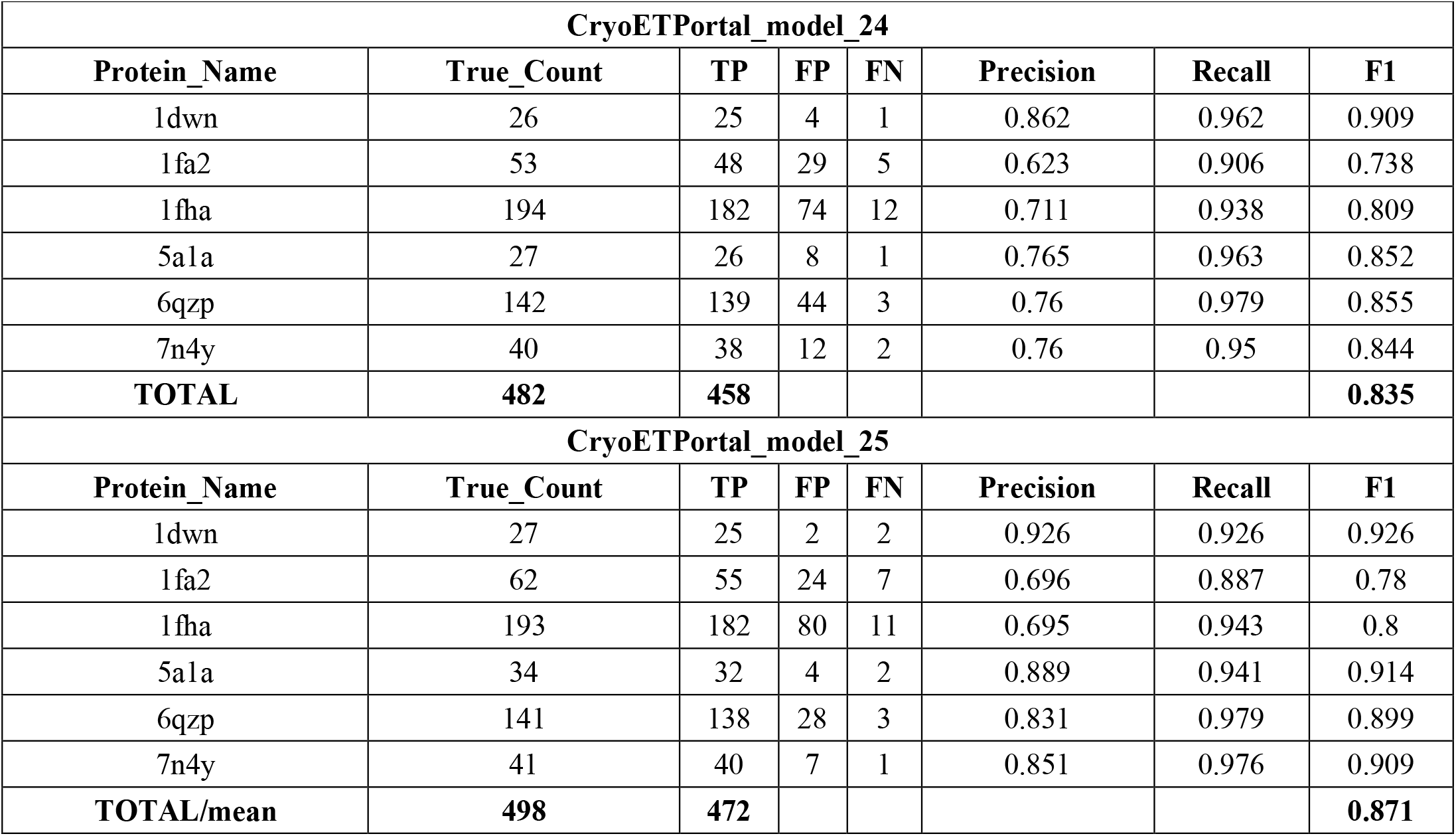
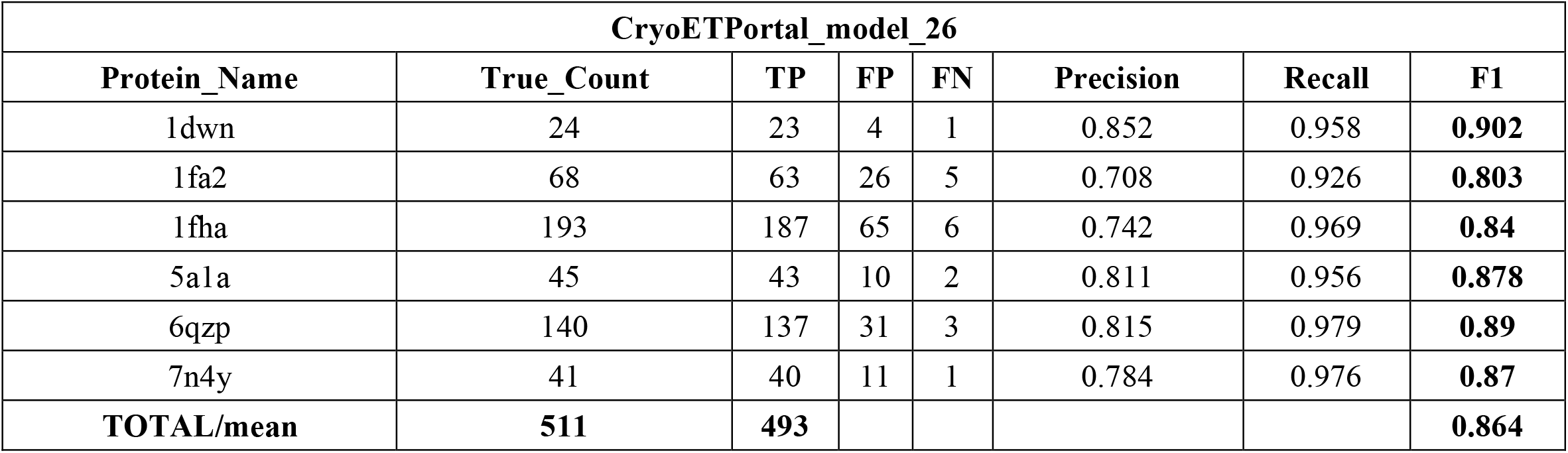
Multiclass particle detection performance metrics of TomoSwin3D on the CryoETPortal test data.

**Supplementary Table S15:**
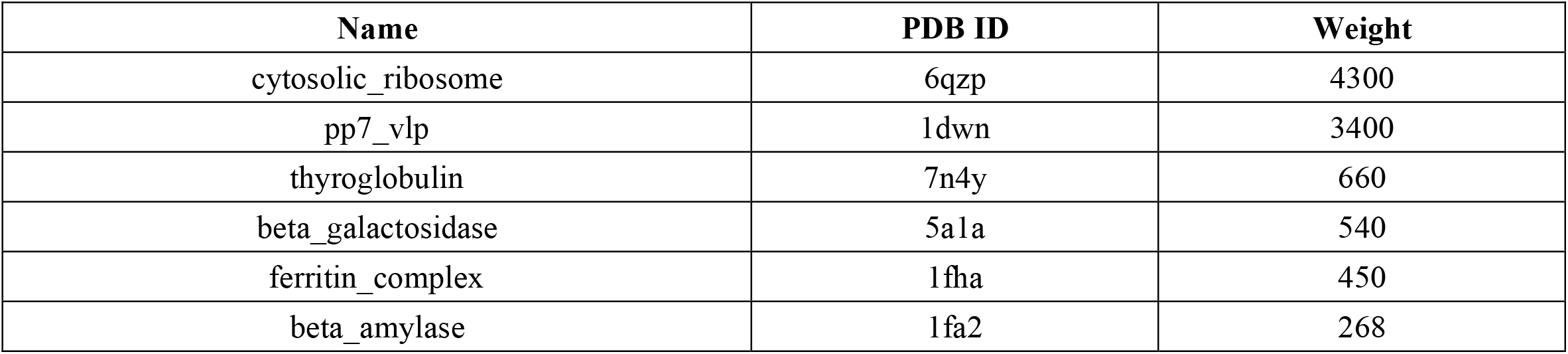
Protein name, corresponding PDB ID and their weight in CryoETPortal dataset.

