## Supplementary information for "TomoSwin3D: a Swin3D Transformer for the Identification and Classification of Macromolecules in 3D Cryo-ET Tomograms"

***Supplementary Table S1:*** *Multiclass particle detection performance metrics of TomoSwin3D on the SHREC 2021 (synthetic) test data*

| Class | GT | TP | FP | FN | Precision | Recall | F1 |
|---|---|---|---|---|---|---|---|
| 1s3x | 122 | 85 | 104 | 37 | 0.45 | 0.697 | 0.547 |
| 3qm1 | 120 | 90 | 110 | 30 | 0.45 | 0.75 | 0.563 |
| 3gl1 | 123 | 92 | 57 | 31 | 0.617 | 0.748 | 0.676 |
| 3h84 | 144 | 132 | 22 | 12 | 0.857 | 0.917 | 0.886 |
| 2cg9 | 125 | 113 | 39 | 12 | 0.743 | 0.904 | 0.816 |
| 3d2f | 140 | 129 | 11 | 11 | 0.921 | 0.921 | 0.921 |
| 1u6g | 143 | 129 | 44 | 14 | 0.746 | 0.902 | 0.816 |
| 3cf3 | 139 | 134 | 2 | 5 | 0.985 | 0.964 | 0.975 |
| 1bxn | 135 | 134 | 3 | 1 | 0.978 | 0.993 | 0.985 |
| 1qvr | 127 | 124 | 2 | 3 | 0.984 | 0.976 | 0.98 |
| 4cr2 | 115 | 112 | 4 | 3 | 0.966 | 0.974 | 0.97 |
| 5mrc | 121 | 120 | 2 | 1 | 0.984 | 0.992 | 0.988 |
| Fiducial | 11 | 11 | 0 | 0 | 1 | 1 | 1 |
| **Macro mean** | | 1405 | 400 | 160 | 0.808 | 0.887 | 0.861 |

***Supplementary Table S2:*** *Details of macromolecular complexes present in the SHREC 2021 dataset, sourced from SHREC 2021 competition.*

| PDB | Name | Mol. weight (kDa) | Volume (nm³) | Area (nm²) | Sphericity | Eff. radius (nm) |
|---|---|---|---|---|---|---|
| 1s3x | Hsp70 ATPase | 42.75 | 90.82 | 109.8 | 0.89 | 2.481 |
| 3qm1 | LJ0536 S106A | 62.62 | 127.9 | 137.6 | 0.892 | 2.789 |
| 3gl1 | Ssb1, Hsp70 | 84.61 | 196.5 | 191.2 | 0.855 | 3.083 |
| 3h84 | GET3 | 158.08 | 347 | 370.9 | 0.644 | 2.807 |
| 2cg9 | Hsp90-Sba1 | 188.73 | 401.2 | 358.4 | 0.734 | 3.358 |
| 3d2f | Sse1p, Hsp70 | 236.11 | 516 | 459.6 | 0.677 | 3.368 |
| 1u6g | Cand1-Cull1-Roc1 | 238.82 | 499.3 | 450.2 | 0.676 | 3.327 |
| 3cf3 | P97/vcp | 541.74 | 1136 | 745.2 | 0.707 | 4.573 |
| 1bxn | Rubisco | 559.96 | 1021 | 583.4 | 0.84 | 5.25 |
| 1qvr | ClpB | 593.36 | 1354 | 1063 | 0.557 | 3.821 |
| 4cr2 | 26S proteasome | 1309.28 | 2675 | 1846 | 0.505 | 4.347 |
| 5mrc | Yeast mito ribosome | 3325.59 | 6372 | 3161 | 0.526 | 6.047 |

***Supplementary Table S3:*** *Protein IDs and their numbers present in the SHREC 2021 dataset, sourced from SHREC 2021 competition*

| Protein | Quantity |
|---|---|
| 1s3x | 122 |
| 3qm1 | 120 |
| 3gl1 | 123 |
| 3h84 | 144 |
| 2cg9 | 125 |
| 3d2f | 140 |
| 1u6g | 143 |
| 3cf3 | 139 |
| 1bxn | 135 |
| 1qvr | 127 |
| 4cr2 | 115 |
| 5mrc | 121 |
| Fiducial | 11 |

***Supplementary Table S4:*** *Macromolecular complexes by their molecular weight in kDa, sourced from SHREC 2021 competition.*

| Group | Weight | Proteins |
|---|---|---|
| Small | <200 | 1s3x, 3qm1, 3gl1, 3h84, 2cg9 |
| Medium | 200-600 | 3d2f, 1u6g, 3cf3, 1bxn, 1qvr |
| Large | 600+ | 4cr2, 5mrc |

***Supplementary Table S5:*** *Multiclass particle detection performance metrics of TomoSwin3D on the SHREC 2020 (synthetic) test data*

| Protein | GT | TP | FP | FN | Precision | Recall | Miss Rate | F1 |
|---|---|---|---|---|---|---|---|---|
| 1s3x | 233 | 205 | 28 | 28 | 0.88 | 0.88 | 0.12 | 0.88 |
| 3qm1 | 241 | 216 | 23 | 25 | 0.904 | 0.896 | 0.104 | 0.9 |
| 3gl1 | 229 | 203 | 24 | 26 | 0.894 | 0.886 | 0.114 | 0.89 |
| 3h84 | 240 | 213 | 20 | 27 | 0.914 | 0.887 | 0.113 | 0.901 |
| 2cg9 | 228 | 220 | 18 | 8 | 0.924 | 0.965 | 0.035 | 0.944 |
| 3d2f | 214 | 205 | 17 | 9 | 0.923 | 0.958 | 0.042 | 0.94 |
| 1u6g | 217 | 208 | 13 | 9 | 0.941 | 0.959 | 0.041 | 0.95 |
| 3cf3 | 238 | 231 | 15 | 7 | 0.939 | 0.971 | 0.029 | 0.955 |
| 1bxn | 245 | 238 | 13 | 7 | 0.948 | 0.971 | 0.029 | 0.96 |
| 1qvr | 226 | 218 | 14 | 8 | 0.94 | 0.965 | 0.035 | 0.952 |
| 4cr2 | 231 | 225 | 10 | 6 | 0.957 | 0.974 | 0.026 | 0.966 |
| 4d8q | 240 | 232 | 12 | 8 | 0.951 | 0.967 | 0.033 | 0.959 |
| | **2782** | **2614** | **207** | **168** | **0.926** | **0.940** | **0.060** | **0.933** |

***Supplementary Table S6:*** *Details of macromolecular complexes present in the SHREC 2020 dataset, sourced from SHREC 2020 competition.*

| PDB | Name | Mol. weight (kDa) | Volume ($nm^3$) | Area ($nm^2$) | Sphericity | Eff. radius (nmx) |
|---|---|---|---|---|---|---|
| 1s3x | Hsp70 ATPase | 42.75 | 104.1 | 122 | 0.877 | 2.56 |
| 3qm1 | LJ0536 S106A | 62.62 | 139.1 | 144.9 | 0.896 | 2.88 |
| 3gl1 | Ssb1, Hsp70 | 84.61 | 207 | 202.6 | 0.835 | 3.065 |
| 3h84 | GET3 | 158.08 | 375.3 | 399 | 0.631 | 2.822 |
| 2cg9 | Hsp90-Sba1 | 188.73 | 394.2 | 380.5 | 0.683 | 3.108 |

| | | | | | | |
|---|---|---|---|---|---|---|
| 3d2f | Sse1p, Hsp70 | 236.11 | 521.9 | 497.9 | 0.63 | 3.145 |
| 1u6g | Cand1-Cul1-Roc1 | 238.82 | 498.5 | 488 | 0.623 | 3.065 |
| 3cf3 | P97/vcp | 541.74 | 1123 | 805.7 | 0.648 | 4.181 |
| 1bxn | Rubisco | 559.96 | 978.9 | 614.4 | 0.776 | 4.78 |
| 1qvr | ClpB | 593.36 | 1255 | 1159 | 0.485 | 3.248 |
| 4cr2 | 26S proteasome | 1309.28 | 3085 | 1971 | 0.52 | 4.696 |
| 4d8q | TRiC/CCT | 1952.74 | 2152 | 1331 | 0.606 | 4.85 |

***Supplementary Table S7:*** *Protein IDs and their numbers present in the SHREC 2020 dataset, sourced from SHREC 2020 competition*

| **protein** | **Quantity** |
|---|---|
| 1s3x | 233 |
| 3qm1 | 241 |
| 3gl1 | 229 |
| 3h84 | 240 |
| 2cg9 | 228 |
| 3d2f | 214 |
| 1u6g | 217 |
| 3cf3 | 238 |
| 1bxn | 245 |
| 1qvr | 226 |
| 4cr2 | 231 |
| 4d8q | 240 |
| **Total** | **2782** |

***Supplementary Table S8:*** *Macromolecular complexes by their molecular weight in kDa, sourced from SHREC 2020 competition.*

| Group | Weight | Proteins |
|---|---|---|
| Small | < 200 | 1s3x, 3qm1, 3gl1, 3h84, 2cg9 |
| Medium | 200 - 600 | 3d2f, 1u6g, 3cf3, 1bxn, 1qvr |
| Large | 600+ | 4cr2, 4d8q |

***Supplementary Table S9:*** *Binary particle detection performance metrics of TomoSwin3D on the EMPIAR-10731 (experimental) test data*

| **SN** | **Tomogram_Name** | **True_Count** | **TP** | **FP** | **FN** | **Precision** | **Recall** | **F1** |
|---|---|---|---|---|---|---|---|---|
| 1 | TS_01 | 353 | 311 | 119 | 42 | 0.723 | 0.881 | 0.794 |
| 2 | TS_02 | 368 | 331 | 112 | 37 | 0.747 | 0.899 | 0.816 |
| 3 | TS_03 | 320 | 243 | 111 | 77 | 0.686 | 0.759 | 0.721 |
| 4 | TS_04 | 331 | 285 | 115 | 46 | 0.712 | 0.861 | 0.78 |
| 5 | TS_05 | 212 | 195 | 70 | 17 | 0.736 | 0.92 | 0.818 |
| 6 | TS_06 | 312 | 262 | 105 | 50 | 0.714 | 0.84 | 0.772 |
| 7 | TS_07 | 305 | 238 | 91 | 67 | 0.723 | 0.78 | 0.751 |
| 8 | TS_08 | 378 | 336 | 113 | 42 | 0.748 | 0.889 | 0.813 |
| 9 | TS_09 | 247 | 204 | 144 | 43 | 0.586 | 0.826 | 0.686 |
| 10 | TS_10 | 308 | 265 | 107 | 43 | 0.712 | 0.86 | 0.779 |

| | | | | | | | | |
|---|---|---|---|---|---|---|---|---|
| 11 | TS_11 | 221 | 201 | 77 | 20 | 0.723 | 0.91 | 0.806 |
| 12 | TS_12 | 344 | 268 | 157 | 76 | 0.631 | 0.779 | 0.697 |
| | Total/Mean | | | | | 0.703 | 0.850 | 0.769 |

***Supplementary Table S10:*** *Binary particle detection performance metrics of CFNPicker on the EMPIAR-10731 (experimental) test data*

| Tomogram | True_Count | TP | FP | FN | Precision | Recall | F1 |
|---|---|---|---|---|---|---|---|
| TS_01 | 353 | 282 | 222 | 71 | 0.56 | 0.799 | 0.658 |
| TS_02 | 368 | 287 | 300 | 81 | 0.489 | 0.78 | 0.601 |
| TS_03 | 320 | 237 | 172 | 83 | 0.579 | 0.741 | 0.65 |
| TS_04 | 331 | 255 | 300 | 76 | 0.459 | 0.77 | 0.576 |
| TS_05 | 212 | 153 | 102 | 59 | 0.6 | 0.722 | 0.655 |
| TS_06 | 312 | 256 | 193 | 56 | 0.57 | 0.821 | 0.673 |
| TS_07 | 305 | 229 | 270 | 76 | 0.459 | 0.751 | 0.57 |
| TS_08 | 378 | 299 | 245 | 79 | 0.55 | 0.791 | 0.649 |
| TS_09 | 247 | 180 | 136 | 67 | 0.57 | 0.729 | 0.639 |
| TS_10 | 308 | 234 | 310 | 74 | 0.43 | 0.76 | 0.549 |
| TS_11 | 221 | 179 | 146 | 42 | 0.551 | 0.81 | 0.656 |
| TS_12 | 344 | 268 | 211 | 76 | 0.559 | 0.779 | 0.651 |
| Total/Mean | | | | | **0.531** | **0.771** | **0.627** |

***Supplementary Table S11:*** *Binary particle detection performance metrics of DeepFinder on the EMPIAR-10731 (experimental) test data*

| SN | Tomogram | True_Count | TP | FP | FN | Precision | Recall | F1 |
|---|---|---|---|---|---|---|---|---|
| 1 | TS_01 | 353 | 268 | 179 | 85 | 0.6 | 0.759 | 0.67 |
| 2 | TS_02 | 368 | 283 | 197 | 85 | 0.59 | 0.769 | 0.667 |
| 3 | TS_03 | 320 | 237 | 158 | 83 | 0.6 | 0.741 | 0.663 |
| 4 | TS_04 | 331 | 252 | 161 | 79 | 0.61 | 0.761 | 0.677 |
| 5 | TS_05 | 212 | 153 | 111 | 59 | 0.58 | 0.722 | 0.643 |
| 6 | TS_06 | 312 | 237 | 152 | 75 | 0.609 | 0.76 | 0.676 |
| 7 | TS_07 | 305 | 229 | 159 | 76 | 0.59 | 0.751 | 0.661 |
| 8 | TS_08 | 378 | 302 | 185 | 76 | 0.62 | 0.799 | 0.698 |
| 9 | TS_09 | 247 | 180 | 136 | 67 | 0.57 | 0.729 | 0.639 |
| 10 | TS_10 | 308 | 237 | 165 | 71 | 0.59 | 0.769 | 0.668 |
| 11 | TS_11 | 221 | 155 | 127 | 66 | 0.55 | 0.701 | 0.616 |
| 12 | TS_12 | 344 | 303 | 109 | 41 | 0.735 | 0.881 | 0.802 |
| | Total/Mean | | | | | 0.604 | 0.762 | 0.673 |

***Supplementary Table S12:*** *Binary particle detection performance metrics of DeepETPicker on the EMPIAR-10731 (experimental) test data*

| SN | Tomogram_Name | True_Count | TP | FP | FN | Precision | Recall | F1 |
|---|---|---|---|---|---|---|---|---|
| 1 | TS_01 | 353 | 284 | 167 | 69 | 0.63 | 0.805 | 0.706 |
| 2 | TS_02 | 368 | 286 | 182 | 82 | 0.611 | 0.777 | 0.684 |
| 3 | TS_03 | 320 | 272 | 104 | 48 | 0.723 | 0.85 | 0.782 |

| | | | | | | | | |
|---|---|---|---|---|---|---|---|---|
| 4 | TS_04 | 331 | 255 | 123 | 76 | 0.675 | 0.77 | 0.719 |
| 5 | TS_05 | 212 | 169 | 66 | 43 | 0.719 | 0.797 | 0.756 |
| 6 | TS_06 | 312 | 239 | 127 | 73 | 0.653 | 0.766 | 0.705 |
| 7 | TS_07 | 305 | 265 | 101 | 40 | 0.724 | 0.869 | 0.79 |
| 8 | TS_08 | 378 | 289 | 122 | 89 | 0.703 | 0.765 | 0.733 |
| 9 | TS_09 | 247 | 205 | 85 | 42 | 0.707 | 0.83 | 0.764 |
| 10 | TS_10 | 308 | 238 | 106 | 70 | 0.692 | 0.773 | 0.73 |
| 11 | TS_11 | 221 | 180 | 127 | 41 | 0.586 | 0.814 | 0.682 |
| 12 | TS_12 | 344 | 287 | 160 | 57 | 0.642 | 0.834 | 0.726 |
| | Total/Mean | | | | | 0.672 | 0.804 | 0.731 |

***Supplementary Table S13:*** *Protein IDs and their numbers present in the EMPIAR-10731 test dataset.*

| SN | Tomogram Name | True Count of Particles |
|---|---|---|
| 1 | TS_01 | 353 |
| 2 | TS_02 | 368 |
| 3 | TS_03 | 320 |
| 4 | TS_04 | 331 |
| 5 | TS_05 | 212 |
| 6 | TS_06 | 312 |
| 7 | TS_07 | 305 |
| 8 | TS_08 | 378 |
| 9 | TS_09 | 247 |
| 10 | TS_10 | 308 |
| 11 | TS_11 | 221 |
| 12 | TS_12 | 344 |

***Supplementary Table S14:*** *Multiclass particle detection performance metrics of TomoSwin3D on the CryoETPortal test data*

| **CryoETPortal_model_24** | | | | | | | |
|---|---|---|---|---|---|---|---|
| **Protein_Name** | **True_Count** | **TP** | **FP** | **FN** | **Precision** | **Recall** | **F1** |
| 1dwn | 26 | 25 | 4 | 1 | 0.862 | 0.962 | 0.909 |
| 1fa2 | 53 | 48 | 29 | 5 | 0.623 | 0.906 | 0.738 |
| 1fha | 194 | 182 | 74 | 12 | 0.711 | 0.938 | 0.809 |
| 5a1a | 27 | 26 | 8 | 1 | 0.765 | 0.963 | 0.852 |
| 6qzp | 142 | 139 | 44 | 3 | 0.76 | 0.979 | 0.855 |
| 7n4y | 40 | 38 | 12 | 2 | 0.76 | 0.95 | 0.844 |
| **TOTAL** | **482** | **458** | | | | | **0.835** |
| **CryoETPortal_model_25** | | | | | | | |
| **Protein_Name** | **True_Count** | **TP** | **FP** | **FN** | **Precision** | **Recall** | **F1** |
| 1dwn | 27 | 25 | 2 | 2 | 0.926 | 0.926 | 0.926 |
| 1fa2 | 62 | 55 | 24 | 7 | 0.696 | 0.887 | 0.78 |
| 1fha | 193 | 182 | 80 | 11 | 0.695 | 0.943 | 0.8 |
| 5a1a | 34 | 32 | 4 | 2 | 0.889 | 0.941 | 0.914 |
| 6qzp | 141 | 138 | 28 | 3 | 0.831 | 0.979 | 0.899 |
| 7n4y | 41 | 40 | 7 | 1 | 0.851 | 0.976 | 0.909 |
| **TOTAL/mean** | **498** | **472** | | | | | **0.871** |

| CryoETPortal_model_26 | | | | | | | |
| --- | --- | --- | --- | --- | --- | --- | --- |
| **Protein_Name** | **True_Count** | **TP** | **FP** | **FN** | **Precision** | **Recall** | **F1** |
| 1dwn | 24 | 23 | 4 | 1 | 0.852 | 0.958 | **0.902** |
| 1fa2 | 68 | 63 | 26 | 5 | 0.708 | 0.926 | **0.803** |
| 1fha | 193 | 187 | 65 | 6 | 0.742 | 0.969 | **0.84** |
| 5a1a | 45 | 43 | 10 | 2 | 0.811 | 0.956 | **0.878** |
| 6qzp | 140 | 137 | 31 | 3 | 0.815 | 0.979 | **0.89** |
| 7n4y | 41 | 40 | 11 | 1 | 0.784 | 0.976 | **0.87** |
| **TOTAL/mean** | **511** | **493** | | | | | **0.864** |

***Supplementary Table S15:*** *Protein name, corresponding PDB ID and their weight in CryoETPortal dataset.*

| Name | PDB ID | Weight |
| --- | --- | --- |
| cytosolic_ribosome | 6qzp | 4300 |
| pp7_vlp | 1dwn | 3400 |
| thyroglobulin | 7n4y | 660 |
| beta_galactosidase | 5a1a | 540 |
| ferritin_complex | 1fha | 450 |
| beta_amylase | 1fa2 | 268 |
